# DNA polymerase kappa stabilized by PTBP2 interacts with MRE11 and promotes genomic instability in leukemia cells

**DOI:** 10.1101/2024.08.18.608437

**Authors:** Shristi Lama, Bibhudev Barik, Sajitha IS, Tannistha Sarkar, Sayantan Chanda, Monalisa Behera, Subhankar Priyadarshi Behera, Sutapa Biswas, Sonali Mohapatra, Ghanashyam Biswas, Soumen Chakraborty

## Abstract

Polypyrimidine tract binding protein 2 (PTBP2) is an RNA-binding protein that controls alternative splicing in neuronal, muscle, and Sertoli cells. Our study unveils a novel role of PTBP2 in promoting the excessive production of the DNA polymerase kappa (*Pol*_κ_) by stabilizing its 3’UTR. We observed an association between its increased expression and the upregulation of PTBP2 in clinical samples of Chronic Myeloid Leukemia (CML). *Ptbp2* knock-out CML cell lines and patient samples treated with hydroxyurea presented with increased DNA damage, as evidenced by long comet tails and higher levels of the DNA damage marker, γH2AX foci, however overexpression of *Pol*_κ_ in the *Ptbp2*-KO cells restored normal phenotype. The deregulation of the DNA repair pathway is a defining feature of malignancies and is closely associated with genomic instability. POLK was found to interact with MRE11 of the MRN complex, thereby governing the activation of ATM-CHK2. Cells with elevated levels of *Ptbp2* and *Pol*_κ_ demonstrated increased sister chromatid exchange and BrdU incorporation in *ex*-*vivo* assays, while multinucleated cells with multipolar spindles were observed in *in*-*vivo* assays. Our findings confirm the critical role of the PTBP2-POLK axis in driving genomic instability and bolstering the viability of cells with increased malignancy.

## Introduction

CML, a myeloproliferative disorder, is characterized by the causative fusion of the Philadelphia chromosome, t (9;22) (q34; q11.2), which results in the BCR::ABL1 chimeric gene with the biological ability, intrinsic or acquired, to cause leukemia (**Chereda et al., 2015**). This fusion leads to the constitutively activated tyrosine kinase activity that triggers several associated downstream pathways, underscoring the severity of the disease (**Ciftciler et al., 2021**). The tyrosine kinase inhibitors have been reported to have excellent clinical benefits for the past 20 years, but not in the blast crisis phase of this inexorably progressing disease (**Hochhaus et al., 2017; Jain et al., 2017**). Advanced molecular characterization revealed clinical diversity and helped to understand drug resistance and CML disease progression. Previously, we reported that BCR::ABL1 potentiates the inclusion of exon 10 of *Ptbp2*, which is essential for its functional protein expression. The inclusion of exon 10 was found in the blast crisis phase of the disease, thereby establishing its role in the progression of CML (**Nandagopalan et al., 2019**). PTBP2 is required for the alternative splicing of precursor mRNA, a process critical for generating multiple protein isoforms from a single gene (**Hannigan et al., 2017**). PTBP2 has also been reported to be highly expressed in cancer cells. It has been proven to promote cell migration and growth of cancer cells (**Ji et al., 2014)**. Recently, we have shown that PTBP2 promotes cell proliferation and tumor formation while enhancing autophagy through Bnip3, thereby supporting the role of PTBP2 as an oncogene in CML (**Barik et al., 2024**). An important feature associated with the disease progression in CML is the accumulation of mutations in the genes that regulate cellular growth and differentiation. Experimental evidence strongly suggests that activated oncogenes that trigger replication stress drive precancerous lesions and cancers. This stress leads to the stalling and collapse of DNA replication forks, forming DNA double-strand breaks (DSBs). The continuous occurrence of DSBs, coupled with error-prone repair processes, could play a crucial role in the genomic instability seen in most human cancers. However, the mechanisms through which these cells acquire genetic alterations and genomic instability remain insufficiently understood. One of the possible mechanisms well reported in the literature is the reactive oxygen species produced by the oncogene BCR::ABL1, which generates endogenous DNA break and the defective repair mechanisms that lead to increased genomic instability, favoring disease progression (**Nowicki et al., 2004; Skorski et al., 2002 and 2007; Sattler et al., 2000)**. The deregulated DNA damage repair pathways have also been reported to be associated with tyrosine kinase inhibitor resistance; **Dinis et al., 2012** have reported the overexpression of base excision repair genes MBD4 and NTLH1 in imatinib-resistant K562 cells. BCR::ABL1 activates the ATR-CHK1 signaling pathway in response to genomic stress, thus establishing a background of the DNA damage and repair in CML disease progression (**Stetka et al., 2020; Shafman et al., 1997)**. Regardless of whether BCR::ABL1 has a direct or an indirect role in promoting genomic instability, 60-80% of patients with CML develop additional non-random chromosomal abnormalities involving chromosomes 8,17, 19, and 22, with duplication of the Ph chromosome or trisomy 8 being the most frequent (**Johansson et al., 2002**). The increasing presence of both γH2AX and 53BP1 foci in cells as a patient transit from the chronic phase to the blast crisis indicates that the promotion of NHEJ and MMEJ serves as a critical mechanism for DNA repair during blastic transformation (**Popp et al., 2020**).

DNA polymerase kappa contributes to the replication checkpoint response and is required for recovery after replication stress. Overexpression of the *Pol*_κ_ has been reported to contribute to aneuploidy, genetic heterozygosity, and DNA breaks, thereby causing genomic instability in different carcinomas (**O-Wang et al., 2001; Bavoux et al., 2005**). It has been proposed that enhanced *Pol*_κ_ expression induces genomic instability by redirecting POLK to the replication forks, thereby interfering with the normal replication process (**Jones et al., 2012**). The overexpression of *Pol*_κ_ causes an increase in the origin of firing, increases the frequency of homologous recombination, and stimulates DNA exchanges and aneuploidy (**Pillaire et al. 2007**). The overexpression of *Pol*_κ_ has been associated with temozolomide resistance, and its inhibition markedly sensitized the cells towards temozolomide as the inhibition of *Pol*_κ_ disrupted the HR-mediated repair and ATR-CHK1 activation (**Peng et al. 2016)**.

In this study, we have unveiled the unique role of an RNA-binding protein, PTBP2, in stabilizing the translesion DNA synthesis enzyme POLK and recruiting the MRN complex via interacting with MRE11 and stimulating the downstream ATM and Chk2 signaling pathway, ultimately fostering disease progression. This axis may be therapeutically targeted to treat the CML blast crisis.

## Materials and methods

### Cell lines and patient samples

The KCL22, K562, KU812, KYO-1 & LAMA84 CML cell lines, as well as the TF1, HEL, HNT34 & F36P AML cell lines, were cultured in RPMI medium 1640 (Gibco, Cat #31800014) supplemented with 10% fetal bovine serum (FBS) (Gibco, Cat. #10270106). The 32Dcl3 cells were cultured in RPMI medium 1640 supplemented with 10% FBS and IL3. The cells were cultured in a humidified incubator at 37°C with 5% CO_2_. Meanwhile, the HEK293T human embryonic kidney cell line was cultured in Dulbecco’s modified Eagle medium (Gibco, Cat. #12100046) and supplemented with 10% FBS. No antibiotic supplements were used unless otherwise specified. All the cell lines were tested for mycoplasma, and mycoplasma was negative. Blood samples were collected from CML patients in the hospital laboratory by venepuncture in K3-EDTA tubes. The institutional human ethical committee approved the study, and the patient’s peripheral blood samples were collected after obtaining written informed consent.

### Generation of KO of Ptbp2 and Pol**_κ_** in cell lines and patient samples and Hydroxyurea treatment

The study used lentivirus-based precision sgRNA Ptbp2 and sgRNA Polκ (Horizon, Cat. # VSGH12606-256402768) to knockout Ptbp2 and Polκ from KCL22 and KU812 cells with puromycin as a selection marker (**Barik et al., 2024 for Ptbp2 KO and the present study for Pol**_κ_ **KO)**. Peripheral blood mononuclear cells (PBMC) were isolated using the Ficol gradient method. Knockout of Ptbp2 was produced in the mononuclear cells by transducing specific sgRNA for Ptbp2 exon 1 (Horizon, Cat. #VSGH11936-247759039). RT-qPCR and Western blotting confirmed Ptbp2 knockout. Cells were treated with Hydroxyurea (HiMedia Cat#H0310) (2mM) for 4h followed by recovery of 12h.

### Generation of stable cell lines overexpressing PTBP2 and POLK

Codon-optimized Ptbp2 cloned in the MSCV-Neo vector was co-transfected with the pCL-Eco vector in the 293T cells to generate the vector control and Ptbp2 retroviral particles. Subsequently, the virus was individually introduced into the murine myeloblastic progenitor cell line, 32Dcl3, and the selection of clones was carried out using neomycin. The study used lentivirus-based precision lentiORF PTBP2 w/stop codon to overexpress Ptbp2 (Horizon, Cat. #OHS5899-202618163) in LAMA84 cells with blasticidin as a selection marker (**Barik et al., 2024)**. The coding region of *Pol*_κ_ was cloned in a lentivirus vector, and the lentivirus was produced by transfection of *Pol*_κ_ plasmid along with packaging plasmid pPAX2 and envelop plasmid pMD2G into HEK293T cells by using standard transfection protocol. Further, *Ptbp2* KO cells were transduced with *Pol*_κ_ lentivirus using polybrene (8µg/ml) and checked for GFP expression. GFP-positive cells were sorted, and POLK expression was checked using Western blot analysis.

### RNA isolation, microarray, cDNA preparation, and RT-qPCR

RNA isolation was done using Trizol (Ambion Cat#15596018) according to the manufacturer’s protocol. The quality of the RNA was checked in the Tape Station and subjected to microarray analysis using the Affymetrix GeneChip Mouse Exon 1.0 ST Array (Catalogue No: 900818) as per the manufacturer’s instruction. RT-qPCR was used with the GoTaq qPCR master mix (Promega Cat # A6002). GAPDH was used as a loading control. The oligos used for qRT-PCR in this study are listed in Supplementary Table 1.

### Western Blotting

Isolation of cell protein lysate and Western blotting was done, as mentioned in **Barik et al., 2024**. The membrane was probed with the following antibodies: GAPDH (CST Cat# 5174S), POLK (Santa Cruz Cat# 166667), PTBP2 (CST Cat# 15719S), MRE11 (CST Cat 4895S), pMRE11 Ser 676 (CST Cat#4859), CHK2 (CST Cat# 2662S), pCHK2 Thr68 (CST Cat#2661), □H2AX Ser 139 (CST Cat#2577). The following secondary conjugated antibodies were used: Anti-rabbit HRP-linked antibody (CST Cat#7074) and Anti-mouse HRP-linked antibody (CST Cat#7076).

### Immunofluorescence

Cells cultured in RPMI with 10% heat-inactivated serum were attached to the coverslips coated with poly-l-lysine (Sigma, Cat# P4707). The coverslips with attached cells were subjected to fixation by Methanol and Acetone in a ratio of 1:1 for 20 min at -20°C. The coverslips were washed with 1x PBS, and blocking was done with 3% BSA (Thermo Fisher Scientific) for 1h. The following antibodies were used: POLK (Santa Cruz, Cat#166667) and γH2AX Ser 139 (CST, Cat#2577). The coverslips were incubated overnight in a moist chamber at 4°C. Secondary antibodies Anti-rabbit 594 Alexa Fluor (Invitrogen, Cat#A11072) and anti-mouse 488 Alexa Fluor (Invitrogen, Cat# A21202) were used and incubated for 1h at dark. The coverslips were stained with DAPI (Merck, Cat#10236276001) for 90 seconds and were mounted in the slides with the mounting media (Invitrogen, Cat #S36938). The slides were visualized using a confocal microscope at 63X (Leica, Germany).

### Alkaline comet assay

Cells were harvested, treated with Hydroxyurea (2mM), and blended with 0.5% low melting agarose. The cells were then added to 1% agarose-precoated slides. The slides were lysed and denatured, followed by electrophoresis. The slides were stained with DAPI and viewed in a fluorescence microscope. The DNA tail percentage was measured using CASP software.

### Actinomycin D chase assay

2.8×10^6^ KCL22, Ptbp2 KO KCL22, LAMA84, Ptbp2 O/E LAMA84 cells were maintained in RPMI containing 10%FBS in 5% CO_2_ incubator at 37 □ Cells were treated with Actinomycin D (Sigma, Cat#A4262) @5ug/ml for different time points: 1h, 2h, 4h, 6h, 8h, and 12h. RNA isolation and cDNA preparation were done using the standard protocol. RT-qPCR was done using GAPDH as a control. The mRNA stability was calculated using GraphPad Prism 7.

### RNA immunoprecipitation (RIP)

RNA immunoprecipitation was carried out as previously described by **Barik et al. 2024**. RT-qPCR was used to quantify the transcript amount.

### Luciferase reporter assay

The 3’UTR of *Polk*, comprising the PTBP2 binding site, was cloned in a luciferase vector, and a luciferase reporter assay was performed as described by **Barik et al. 2024**. The binding site was mutated using site-directed mutagenesis.

### Cell survival assay

Cells (5000/well) were seeded in a 24-well plate. After treatment with Hydroxyurea 2mM for different time points, cells were allowed to grow for **7** days. Cell survival was monitored by MTT (Sigma, #M5655, 0.1mg/ml) assay and reading taken at 570 nm in VICTOR Nivo^TM^ multimode reader (PerkinElmer, USA). Percent survival was calculated as NTC cells/ Ptbp2 KO cells*100.

### BrdU incorporation assay

100uM of bromodeoxyuridine (Sigma, Cat#B5002) was incorporated in the DNA for 20 min to visualize the replication foci. Cells were fixed with 70% cold ethanol. Cells were denatured in 1.5M HCL for 30 min and incubated in 5% BSA, 0.5% Tween 20, and 5% FBS for 20 min. Primary Anti-Brdu (Merck, Cat#B2531) antibody was added to the slides precoated with poly-l lysine (Sigma, cat# P4707) and incubated for 1h. Slides were washed with PBS-Tween20 and incubated with a secondary antibody (Invitrogen, Cat#A21202) for 1h.

### Metaphase spreads

1×10^6^ cells were seeded in RPMI+10%FBS in a humidified chamber at 5% CO_2_ chamber at 37°C. Cells were treated with colchicine (Sigma, Cat#C9754) for 45 min before harvesting. Cells were treated with KCL 0.056M solution for 30 min and fixed with methanol: glacial acetic acid (3:1) solution. The cell suspension was taken in a Pasteur pipette, and a single drop was released from a height onto the slide. The slides were allowed to air dry. The slides were stained with Giemsa (Himedia, Cat#TCL083) for 30 min. The images were acquired in an Apotome microscope (Zeiss).

### DNA Fiber assay

The cells were labeled with 100uM of CIdu (Sigma, Cat#C6891) and 100uM of IdU (Sigma, Cat#I712) for the indicated times. HU was added, as shown in the result. Cells were harvested, lysed, and stretched in super-frosted slides. Cells were denatured with 2.5M HCL and blocked with a blocking solution. The slides were incubated with primary antibodies, Rat anti-CldU 1:100 (Abcam, Cat#ab6326) and Mouse Anti-IdU 1:100 (Becton Dickinson, Cat# 347580) for 2.5 hours at RT, washed with PBS and incubated with Rat anti-Cy3 (Jackson Immuno Research, Cat#712-166-153), Mouse Alexa 488 (Molecular probes, Cat#A11001) secondary antibodies. Following mounting, slides were imaged using a confocal microscope at 63X (Leica, Germany).

### Animal study

All animal protocols were performed with the approval of the Institute of Life Sciences ethics committee (ILS/IAEC-03-AH/05-14 and ILS/IAEC-159-AH/AUG-19). The subcutaneous tumor model mentioned in Barik et al. 2024 and the transplantation model, B6-CD45.1 and CD45.2 mice, were used per the standard protocol. All mice were bred in the animal house facility. Lineage-negative cells were isolated from 8-10 weeks CD45.1 mice with a lineage-negative cell isolation kit (Miltenyi Biotech, Cat#130-110-470) and transduced with MIGR1 vector, BCR-ABL and BCR-ABL+Ptbp2 virus. Transduced bone marrow cells were transplanted into 8-10 weeks lethally irradiated (10 Gy) CD45.2 mice. Control mice were transplanted with MIGR1vector transduced lineage-negative cells. After 30 days, the mice were sacrificed, and the liver, kidney, and spleen were isolated and stored in formalin.

### Immunohistochemistry (IHC)

IHC staining for the tissue was performed using the protocol described previously (**Barik et al., 2024**). Each slide marked one of the sections as a negative control, on which the primary antibody was omitted. Images were obtained using a Leica DM500 microscope (Leica, Germany). Antibodies used for the assay: Polk (Santa Cruz, Cat# 166667), PTBP2 (CST, #Cat 15719), γH2AX Ser 139 (CST, #2577).

### H&E staining

Formalin-fixed and paraffin-embedded 4-5μm thick sections were taken. The sections fixed in the slides were washed three times with Xylene (Himedia, Cat# AS080), followed by dehydration with 100% ethanol, 95% ethanol, 80% ethanol, and 70% ethanol each for 5 min. The slides were stained with Haematoxylin (Himedia, Cat# S058) for 2 min and washed in tap water. The slides were stained with Eosin (Himedia, Cat#3007), rehydrated with 90% and 100% ethanol for 30 seconds, and xylene for 30 seconds. The slides were mounted in DPX (Merck, Cat#06522), followed by imaging.

### Statistical analysis

GraphPad Prism 7.0 was used to analyze the data and create the graphs. Statistical analysis was performed using two-way ANOVA for grouped analysis or paired *t*-test for 2 groups. All data are represented as the mean ± SEM. *, p < 0.05, * *, p < 0.01 and * **, p < 0.001 were considered as statistically significant. All experiments were performed at least thrice with three biological repeats.

## Results

### PTBP2 targets the 3’-UTR of DNA polymerase kappa and stabilizes its expression

Our recent findings have revealed a significant overexpression of PTBP2 in Chronic Myeloid Leukemia (CML) and Acute Myeloid Leukemia (AML) cell lines. This overexpression has been shown to promote cell proliferation by facilitating mitochondrial fusion and enhancing autophagy through the upregulation of BNIP3 (**Barik et al., 2024**). To find the targets of PTBP2 in the hematopoietic cells, we overexpressed *Ptbp2* in the 32Dcl3 cells, a murine myeloid progenitor cell line. RT-qPCR showed expression of *Ptbp2* in *Ptbp2*-32Dcl3 cells compared to the vector-32Dcl3 cells (**Fig. 1A, left panel**). Western blot confirmed the expression of PTBP2 in the 32Dcl3 cells (**Fig. 1A, right panel**). Gene expression was performed in the vector-32Dcl3 and *Ptbp2*-32Dcl3 cells using the Affymetrix GeneChip Mouse Exon 1.0 ST Array. A subset of the statistically significant gene set, with the cutoffs of Log2 fold change range of ≤2.0 and ≥-2.0 and p-value <0.05, were taken into consideration and represented in the heat map (**Supplementary Fig. 1A**). Differential gene expression of the targets was verified using RT-qPCR (**Fig. 1B**). The relative expression of DNA polymerase kappa (*Pol*_κ_) was found to be highest in the presence of PTBP2. The expression of PTBP2 and POLK in protein levels was checked in different CML and AML cell lines. The expression of PTBP2 was identical in all the cell lines, as reported earlier (**Barik et al., 2024**). The expression of POLK was found to be similar to PTBP2 in the CML and AML cell lines (**Fig. 1C**). As reported earlier, *Ptbp2*-KO-KCL22, *Ptbp2*-KO-KU812, and the corresponding NTC cells were used in the study. The methodology for the generation of stable *Ptbp2*-KO cell lines is outlined in **Barik et al., 2024**. RT-qPCR and western blot analysis confirmed the knockout of *Ptbp2* from the cells (**Barik et al., 2024**). Reduced expression of *Pol*_κ_ was observed in the *Ptbp2*-KO-KCL22 and *Ptbp2*-KO-KU812 with respect to the NTC cells at the mRNA (**Supplementary Fig. 1B**) and the protein level (**Fig. 1D, upper panel**). Overexpression of *Ptbp2* in the CML cell line LAMA84 was also reported earlier (**Barik et al., 2024**). In this study, we found upregulation of *Pol*_κ_ in the *PTBP2*-O/E-LAMA84 cells with respect to the vector-transduced LAMA84 cells both at the mRNA (**Supplementary Fig. 1C)** and the protein level (**Fig. 1E**).

**Figure 1.**
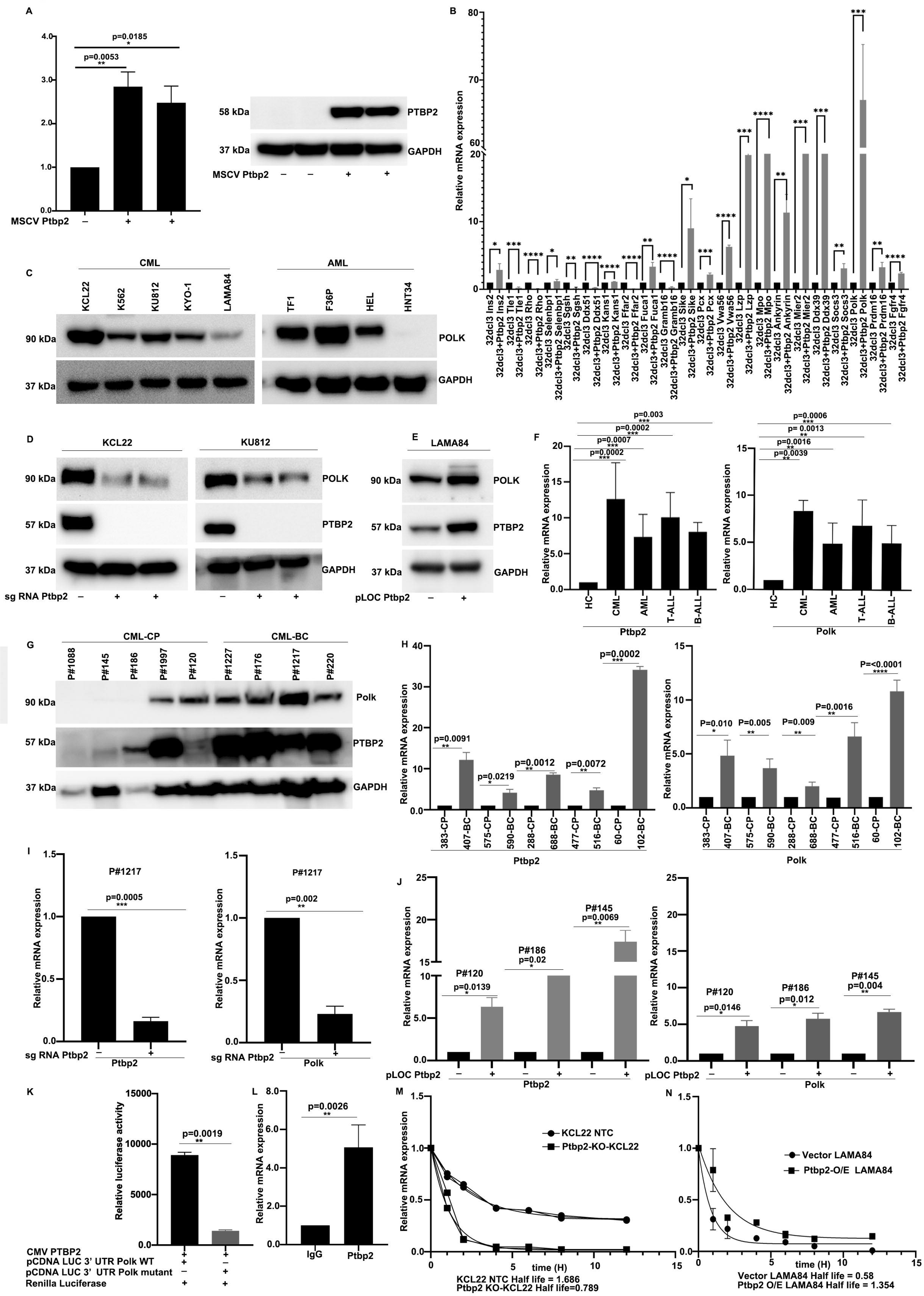
PTBP2 targets DNA polymerase kappa and facilitates its upregulation. A) mRNA and protein expression of PTBP2 in vector-32Dcl3 and 32Dcl3-Ptbp2 cells. B) Differential expression of the targets by RT-qPCR. C) Western blot analysis for POLK and GAPDH in different CML and AML cell lines. D) Western blot analysis of POLK (upper panel) and PTBP2 (middle panel) in WT NTC cells and two different clones of Ptbp2 ^-/-^ in each of the KCL22 and KU812 cells. GAPDH was used as a loading control. E) Expression of POLK in Ptbp2 overexpressed LAMA84 cells. F) Relative mRNA expression of Polκ and Ptbp2 in healthy control (HC), CML, AML, T-ALL, and B-ALL patient samples. G) Expression of POLK in 5 CML CP samples and 4 CML BC samples. H) RT-qPCR analysis of Ptbp2 and Polκ in 5 paired CML patient samples. I) Genetic ablation of Ptbp2 in CML BC sample and RT-qPCR analysis of Ptbp2 and Polκ respectively ***, P=0.0005, **, P =0.002. J) Overexpression of Ptbp2 in CML CP samples and RT-qPCR analysis of Ptbp2 and Polκ, respectively. K) HEK293T cells were co-transfected with CMV Ptbp2, pCDNA LUC 3’UTR Polκ WT, and pCDNA LUC 3’UTR Polκ mutant. Renilla luciferase was used as an internal control for 48 h. The results were expressed as mean ± SEM of triplicate experimental analysis. L) RT-qPCR of PTBP2 bound Polκ transcript with respect to IgG control. M) mRNA half-life of Polκ transcript in KCL22 NTC and Ptbp2-KO-KCL22 cells. N) mRNA half-life of Polκ transcript in LAMA84 and Ptbp2 O/E LAMA84 cells.

Next, we checked the expression of *Ptbp2* and *Pol*_κ_ in both myeloid (CML and AML) and non-myeloid (B-ALL and T-ALL) samples by RT-qPCR. 34 CML, 12 AML, 11 T-ALL, 11 B-ALL samples, and 5 healthy control (HC) samples were considered. The expression of *Ptbp2* and *Pol*_κ_ were higher in the samples than in the healthy controls (HC) (**Fig. 1F**). The correlation between the two genes was positive in CML and AML samples (**Supplementary Fig. 1D)**. We checked for the protein expression of PTBP2 and POLK in the PBMC of CML CP (P#1088, P#145, P#186, P#1997, P#120) and CML BC samples (P#1227, P#176, P#1217, P#220) (**Fig 1G**). CML CP samples P#1088 and P#145 did not show any expression of PTBP2 and POLK, P#186 expressed only PTBP2 but not POLK, P#1997 expressed both PTBP2 and POLK whereas P#120 showed only the expression of POLK, but not PTBP2. CML BC samples (P#1227, P#176, P#1217, and P#220) showed expression of both PTBP2 and POLK. Earlier, we have demonstrated that the expression of *Ptbp2* increases with the progression of the disease (**Nandagopalan et al., 2019**). Our data suggest that the expression of PTBP2 and POLK is correlated in CML BC samples, and the variation in the expression in the CML CP sample may be due to interpatient heterogeneity (**Fig. 1G**). The expression of *Ptbp2* and *Pol*_κ_ was also checked in five paired (CP to BC) CML cases. Both *Ptbp2* and *Pol*_κ_ were found to be upregulated with the progression of the disease (**Fig. 1H**). Again, as the CML BC sample (P#1217) was observed to have a very high expression of *Ptbp2* both by RT-qPCR and western blot and the PBMC was available with us, we knocked out *Ptbp2* in these cells using specific sgRNA for *Ptbp2*. Expression of *Ptbp2* and *Pol*_κ_ was found to be decreased in the cells (**Fig. 1I**). Likewise, overexpression of *Ptbp2* was done using precision lentiORF *Ptbp2* w/stop codon in the CML CP patient samples (P#120, P#186, P#145) which was found to have a very low expression of *Ptbp2*. Overexpression of *Ptbp2* in these samples elevated the expression of *Pol*_κ_ (**Fig. 1J**).

As the knockout of *Ptbp2* was directly correlated with the decreased expression of *Pol*_κ_, we speculated that PTBP2 may be controlling the expression of *Pol*_κ_. It was reported that PTBP2 binds to the 3’UTR of several genes in the murine male germ cells **(Xu et al., 2007 and 2008).** We examined PTBP2 binding sites on *Pol*_κ_ mRNA by scanning the total *Pol*_κ_ mRNA using beRBP (https://bioinfo.vanderbilt.edu/beRBP/predict.html). Two specific PTBP2 binding sites (CUUUUCU and CUUUCU) were observed in the 3’UTR with binding scores of 0.566 and 0.492, respectively (**Supplementary Fig. 1E**). To confirm, we cloned both binding sites in a luciferase vector. The wild-type *Ptbp2*, the luciferase reporter with the binding site, and the Renilla luciferase were transfected in the 293T cells. An increase in the expression of luciferase reporter with binding site 1 (CUUUUCU) but not site 2 was observed compared to empty vector control. Modification of the binding site 1 (CAAAACA) using site-directed mutagenesis decreased the relative luciferase reporter count, providing evidence that PTBP2 binds and regulates the stability of *Pol*_κ_ (**Fig. 1K**). Again, PTBP2 and its associated RNA were immunoprecipitated with a PTBP2-specific antibody, and the immunoprecipitated PTBP2 was initially checked with the western blot (**Supplementary Fig. 1F**). Subsequently, the binding efficiency of PTBP2 to the associated *Pol*_κ_ mRNA was observed by using RT-qPCR (**Fig. 1L**). The binding of PTBP2 to *Pol*_κ_ was reported earlier, wherein it was observed that PTBP2 is involved in the binding and splicing of *Pol*_κ_ in mouse cells at two different stages, spermatogonia to spermatocyte and spermatocyte to round spermatids; however, no splicing was observed in the case of round spermatids to spermatozoa (**Hannigan et al., 2017**). Again, we used actinomycin D to block transcription and confirm that PTBP2, not unintended effects, directly caused decreased *Pol*_κ_ mRNA. NTC-KCL22 and *Ptbp2*-KO-KCL22 @ 0.5 million cells were treated with 5□μg/mL of actinomycin D, and the RNA was harvested at 0-, 5-, 10-, and 15-h post-treatment. We measured the mRNA half-life of *Pol*_κ_ by determination of transcript abundance by RT-qPCR. Half-life was 2.39h at the *Pol*_κ_ gene in NTC KCL22 cells. In comparison, *Ptbp2*-KO-KCL22 cells exhibited a half-life of 0.84h, indicating a significant decrease in the stability of *Pol*_κ_ in the *Ptbp2* depleted state (**Fig. 1M**). Overexpression of *Ptbp2* in the LAMA84 cells showed a half-life of *Pol*_κ_ of 1.354h with respect to vector-transduced LAMA84 cells, which was 0.58h (**Fig. 1N**). Our results, thus, show that PTBP2 binds to the 3’UTR and regulates the stability of *Pol*_κ_.

### PTBP2-POLK axis acts as a regulator of DNA repair in hematological malignancies

As POLK is involved in error-prone DNA repair, we aimed to investigate the connection between PTBP2 and POLK in DNA repair. Hydroxyurea (HU), a potent inducer of replication stress that promotes DNA damage□independent replication fork arrest, was used (**Lopes et al., 2001; Sogo et al., 2002**). HU is commonly used in cases of CML to lower the high WBC count before starting imatinib. HU initially resulted in stalled replication forks; however, after prolonged treatment, it collapses into DSBs (**Saintigny et al., 2001**). POLK has been reported to play a critical role in fork restart during high-dose HU (2 mM) treatment (**Tonzi et al., 2018**). KCL22-NTC and KU812-NTC and the *Ptbp2*-KO-KCL22 and *Ptbp2*-KO-KU812 cells were treated with HU at a concentration of 2mM for 4 h, followed by a recovery period of 12 h, after which Comet assay was done. Comet assay measures DNA breaks in eukaryotic cells, and the tail length of the comet is directly proportional to the extent of DNA damage in the cells. After HU treatment, the comet tail increased in the *Ptbp2*-KO-KCL22 cells with respect to the KCL22-NTC cells (**Fig. 2A, middle panel)**. We also observed a significant increase in the comet tail, with HU treatment, in the *Ptbp2*-KO-KU812 cells with respect to the KU812-NTC cells and vector-LAMA84 cells with respect to the *Ptbp2*-O/E-LAMA84 cells (**Supplementary Fig. 2A, A and C**). The bar diagram represents the change (**Supplementary Fig. 2A, B, and D**). As *Ptbp2* knockout depletes *Pol*_κ_ and POLK is reportedly involved in the DNA damage response, we overexpressed *Pol*_κ_ in the *Ptbp2*-KO-KCL22 and generated the *Ptbp2*-KO-KCL22-*Pol*_κ_-O/E cells (**Fig. 2B**). When the *Ptbp2*-KO-KCL22-*Pol*_κ_-O/E cells were subjected to comet assay after treatment with HU, they exhibited a small comet tail, almost identical to KCL22-NTC cells (**Fig. 2A, lower panel**). The bar diagram (**Fig. 2C**) represents the change in the comet tail. This demonstrates the significance of the PTBP2-POLK axis in the DNA damage tolerance pathway.

**Figure 2.**
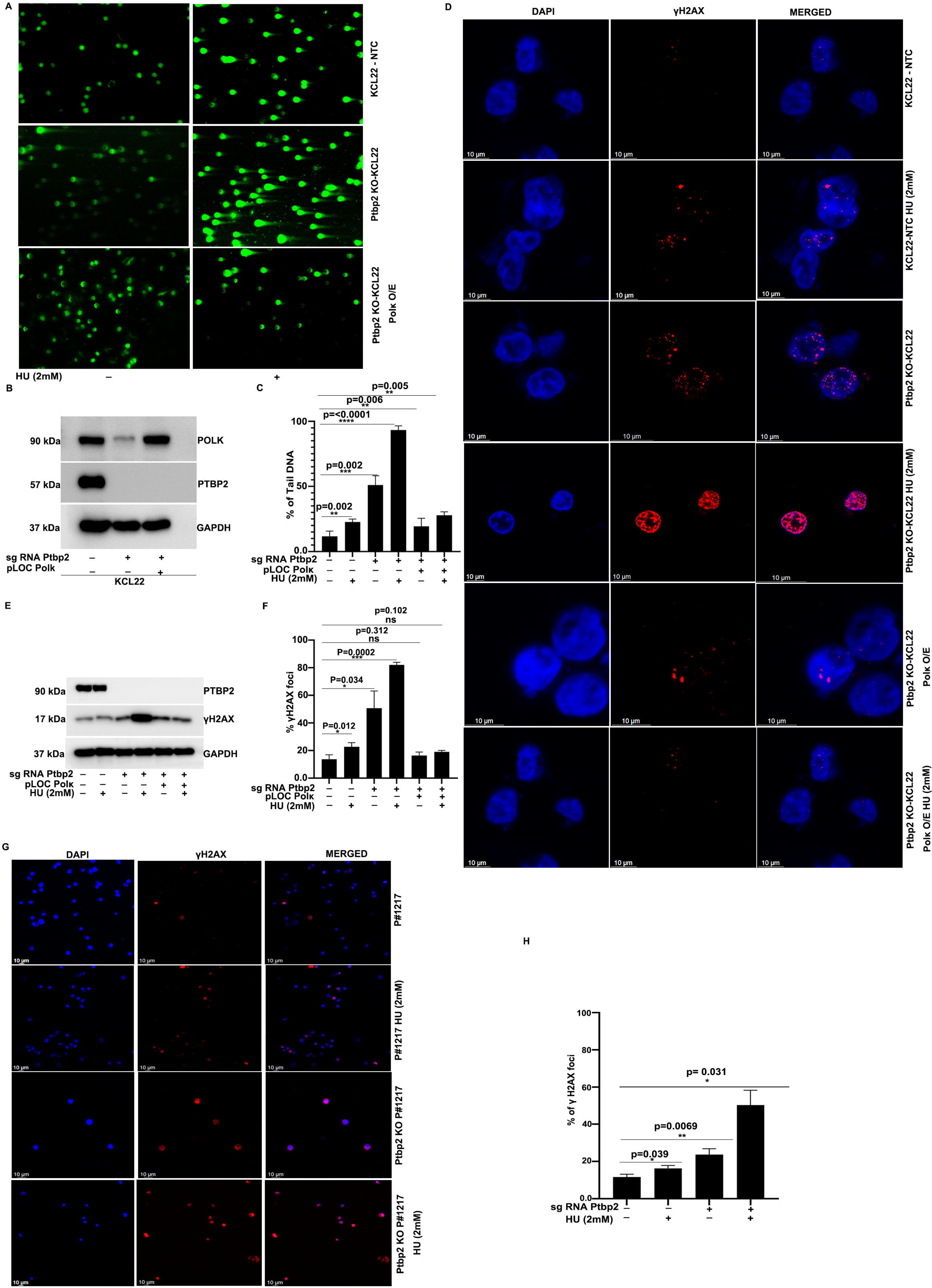
PTBP2-POLK axis acts as a regulator of DNA repair in hematological malignancies. A) KCL22-NTC, Ptbp2-KO-KCL22, and Ptbp2-KO-KCL22-Polκ-O/E cells were treated with or without hydroxyurea and subjected to alkaline comet assay. B) Represents the protein expression of the POLK in the respective cells used for the alkaline comet assay. C) Quantification of the percentage of DNA tail. The data are presented as the mean ± SEM. D) KCL22-NTC, Ptbp2-KO-KCL22, and Ptbp2-KO-KCL22-Polκ-O/E cells treated with or without hydroxyurea and probed with γH2AX antibody and DAPI. E) Protein expression of γH2AX and PTBP2 in KCL22-NTC, Ptbp2-KO-KCL22, and Ptbp2-KO-KCL22-Polκ-O/E cells treated with or without hydroxyurea F) Quantitative analysis of the percentage of γH2AX foci. A cell containing at least 10 foci was considered a foci-positive cell. G) CML BC sample 1217 and Ptbp2-KO-1217 cells were treated with or without hydroxyurea and probed with γH2AX antibody and DAPI. H) Quantitative analysis of the percentage of γH2AX foci. A cell containing at least 10 foci was considered a foci-positive cell.

Phosphorylation of the Ser-139 residue of the histone variant H2AX, forming γH2AX, is an early cellular response to the induction of DSBs (**Bonner et al., 2008**). Again, we checked the cells for the γH2AX foci through immunofluorescence and expression by western blot. The γH2AX foci were higher in the *Ptbp2*-KO-KCL22 than in the KCL22-NTC cells when treated with HU (**Fig. 2D, 4^th^ panel compared to the 2^nd^ panel**). However, little change in the γH2AX foci was observed in the *Ptbp2*-KO-KCL22-*Pol*_κ_-O/E cells with or without HU treatment (**Fig. 2D, 5th, and 6th panel, respectively).** The percentage of γH2AX foci is represented in the graph (**Fig. 2F**). The *Ptbp2*-KO-KCL22 cells, following the γH2AX foci, showed a higher expression of γH2AX in western blot as compared to the KCL22-NTC when treated with HU (**Fig. 2E, middle panel, 4^th^ lane**); however, no change was observed when *Pol*_κ_ was overexpressed in the KO cells **(Fig. 2F, middle panel, 6^th^ lane)**. The γH2AX foci were higher in vector-LAMA84 with respect to the *Ptbp2*-O/E-LAMA84 cells when treated with or without HU (**Supplementary Fig. 2A, E**). The percentage of γH2AX foci and the protein expression are represented respectively (Supplementary Fig. 2A, F, **and G)**. Similar observation was found in KU812-NTC and *Ptbp2*-KO-KU812 cells when treated with HU as the γH2AX foci were higher in the *Ptbp2*-KO-KU812 than in the KU812-NTC cells when treated with HU (**Supplementary Fig 2B, A**) The percentage of γH2AX foci is represented in the graph (**Supplementary Fig 2B, B**) The *Ptbp2*-KO-KU812 cells, following the γH2AX foci, showed a higher expression of γH2AX in western blot as compared to the KU812-NTC cells when treated with HU (**Supplementary Fig 2B, C**). Peripheral blood mononuclear cells isolated from the CML BC sample (P#1217) were treated with or without HU (2 mM) for 4 h, followed by a recovery of 12 h. Here, the expression of γH2AX foci was found to be significantly higher in the *Ptbp2* KO (P#1217) cells than in the NTC cells (P#1217) (**Fig. 2G, last panel**) upon HU treatment. The percentage of γH2AX foci is represented in the graph (**Fig. 2H**). The immunofluorescence and western blot data indicate the presence of DNA damage and impaired repair in the cells lacking *Ptbp2*. Again, we tested whether ablation of PTBP2 leads to a change in cell viability. We treated KCL22-NTC and KU812-NTC, as well as *Ptbp2*-KO-KCL22 and *Ptbp2*-KO-KU812 cells, along with vector-LAMA84 and *Ptbp2*-OE-LAMA84 cells with HU for 2, 4, 6, 8, and 12 h. Fresh media without HU was added after the mentioned time, and the cells were allowed to grow for 6 days. Cell viability was checked using the MTT assay. An almost 50% decrease in cell viability was observed in the *Ptbp2*-KO-KCL22, *Ptbp2*-KO-KU812, and the vector-LAMA84 cells (**Supplementary Fig. 2B-D, E, and F**). Again, the same cells were checked for the percentage of apoptosis. After the HU treatment, increased apoptosis was observed when *Ptbp2* was ablated from KCL22 and KU812 cells and in vector LAMA84 cells (**Supplementary Fig 2C**). All these data suggest that the PTBP2-POLK axis activates the DNA repair pathway and protects the cells from apoptosis at the expense of genomic instability after HU treatment.

### PTBP2 regulates the MRE11-ATM-CHK2 axis via DNA polymerase kappa

MRE11-RAD50-NBS1 are the first sensors for the double-strand break, replication fork collapse, and dysfunction of telomeres **(Nowicki et al., 2004)**. MRE11 acts as a primary sensor and recruits ATM, and its phosphorylation facilitates the activation of CHK2 **(Koptyra et al., 2006)**. We assessed the effect of *Ptbp2*-KO on the MRE11 and its phosphorylation. The expression of MRE11 and phosphorylation significantly decreased in the *Ptbp2-*KO cells compared to its non-targeted control in KCL22 and KU812 cells (**Fig. 3A**). Similarly, the overexpression of *Ptbp2* in LAMA84 cells was positively correlated with the expression of MRE11 and pMRE11 (**Fig. 3B**). Thus, we sought to examine the downstream regulation of MRE11 by PTBP2. We performed the co-immunoprecipitation of PTBP2 to check for the interaction with MRE11; however, we did not observe any interaction between PTBP2 and MRE11(**Fig not shown**). To determine whether POLK affects MRE11 expression, we checked the expression of MRE11 in *Ptbp2* ablated KCL22 cells overexpressed with *Pol*_κ_. The expression of MRE11 and phospho-MRE11 was rescued when *Pol*_κ_ was overexpressed (**Fig. 3C**). To further confirm the regulation of MRE11 by POLK, we genetically ablated *Pol*_κ_ from the KCL22 cells. Then, we checked for the expression of MRE11 and its phosphorylation. A decrease in the expression of MRE11 was observed and its phosphorylation, thus confirming its regulation via POLK (**Fig. 3D**). To examine whether MRE11 is regulated by POLK, we immunoprecipitated POLK and assessed the co-immunoprecipitation for MRE11. MRE11 co-immunoprecipitated with POLK (**Fig. 3E, left panel)**. To further validate the association between POLK and MRE11, we did a reverse co-immunoprecipitation wherein MRE11 was pulled down, and the expression of POLK was checked. The POLK co-immunoprecipitated with MRE11 (**Fig. 3E, right panel**). This data was further validated as POLK and MRE11 were found to colocalize in the CML cell lines (**Fig. 3F**).

**Figure 3.**
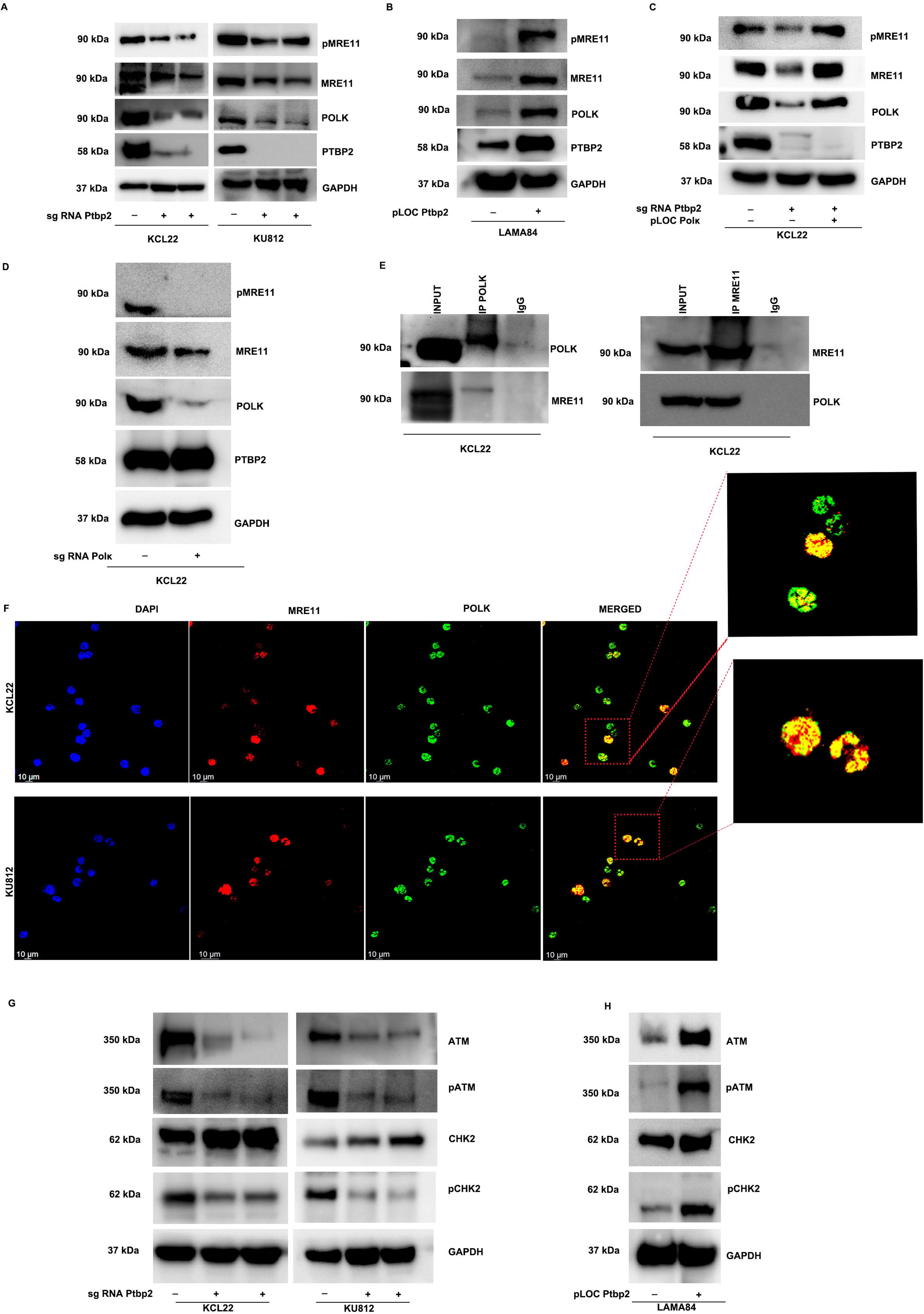
PTBP2 regulates the MRE11-ATM-CHK2 axis. A) MRE11, pMRE11, and POLK were expressed in WT NTC and two Ptbp2-KO-KCL22 and KU812 cell clones. B) Expression of MRE11, pMRE11, and POLK in LAMA84 and LAMA84 O/E Ptbp2 cells. C) MRE11, pMRE11, and POLK expression in WT NTC, Ptbp2-KO-KCL22, and Ptbp2-KO-Polκ-O/E-KCL22 cells. D) Expression of MRE11, pMRE11 in Polκ ablated KCL22 cells. E) Western blot analysis of co-immunoprecipitation of POLK and MRE11. F) Representative images of cells co-immunofluorescence probed with Anti-MRE11 (red) and Anti-POLK (green) are shown in KCL22 and KU812 cells. G) Protein expression of ATM, pATM, CHK2, and pCHK2 in WT NTC and two different Ptbp2-KO-KCL22 and KU812 cell clones. H) Protein expression of ATM, pATM, CHK2, and pCHK2 in LAMA84 and Ptbp2-O/E-LAMA84 cells.

Since MRE11 is the major player in the recruitment of ATM in the DNA damage sites and ATM-CHK2 is an important regulator of DNA damage that protects the cells from DNA damage-induced cytotoxicity, we checked the expression of ATM-CHK2. We observed that the phosphorylation of ATM-CHK2 in KCL22 and KU812 cells was significantly lower in the *Ptbp2*-KO cells than in its non-targeted control cells (**Fig. 3G**). Ectopic expression of *Ptbp2* displayed significantly higher levels of ATM-CHK2 phosphorylation in the LAMA84 cells (**Fig. 3H**). Our data suggests PTBP2 stabilizes the 3’UTR of *Pol*_κ_ in CML-BC cases, facilitating the interaction of POLK with MRE11, thereby promoting error-prone DNA repair. Similarly, the ablation of *Ptbp2* is associated with the destabilization of Polκ, followed by the inactivation of the ATM-CHK2 pathway.

### PTBP2 protects the stalled replication forks from degradation and promotes genomic instability and chromosomal aberration

To investigate the role of PTBP2 in replication fork stabilization, we employed the DNA fiber assay where the NTC and *Ptbp2* ablated KCL22 cells were sequentially labeled with CIdU (red) followed by IdU (green), after which the active forks were stalled with HU (**Fig. 4A)**. The relative shortening of IdU tract after HU treatment serves as the measure of replication fork degradation. Wild-type cells showed a mean IdU/CIdU ratio close to 1 upon HU treatment. However, *Ptbp2*-deficient cells exhibited a 35%-45% reduction in the IdU tract length. The IdU/CIdU ratio was less than 1, depicting that in the absence of *Ptbp2*, the replication forks were not protected from degradation, more so after HU treatment (**Fig. 4B**). Consistent with our data, the replication forks were protected from degradation in *Ptbp2* O/E LAMA84 cells as the IdU/CIdU ratio was close to 1. In contrast, the normal LAMA84 cells underwent replication fork degradation upon HU treatment (**Fig. 4C**). To validate further that *Ptbp2* mediates the replication fork stability via POLK, we performed the sequential labeling of CIdU and Idu followed by HU treatment in *Ptbp2* ablated KCL22 cells overexpressed with *Pol*_κ_. The data suggested that the replication fork in *Ptbp2* ablated KCL22 cells overexpressed with *Pol*_κ_ was stable compared to the *Ptbp2* ablated KCL22 cells, as depicted by the IdU tract length (**Fig. 4D**).

**Figure 4.**
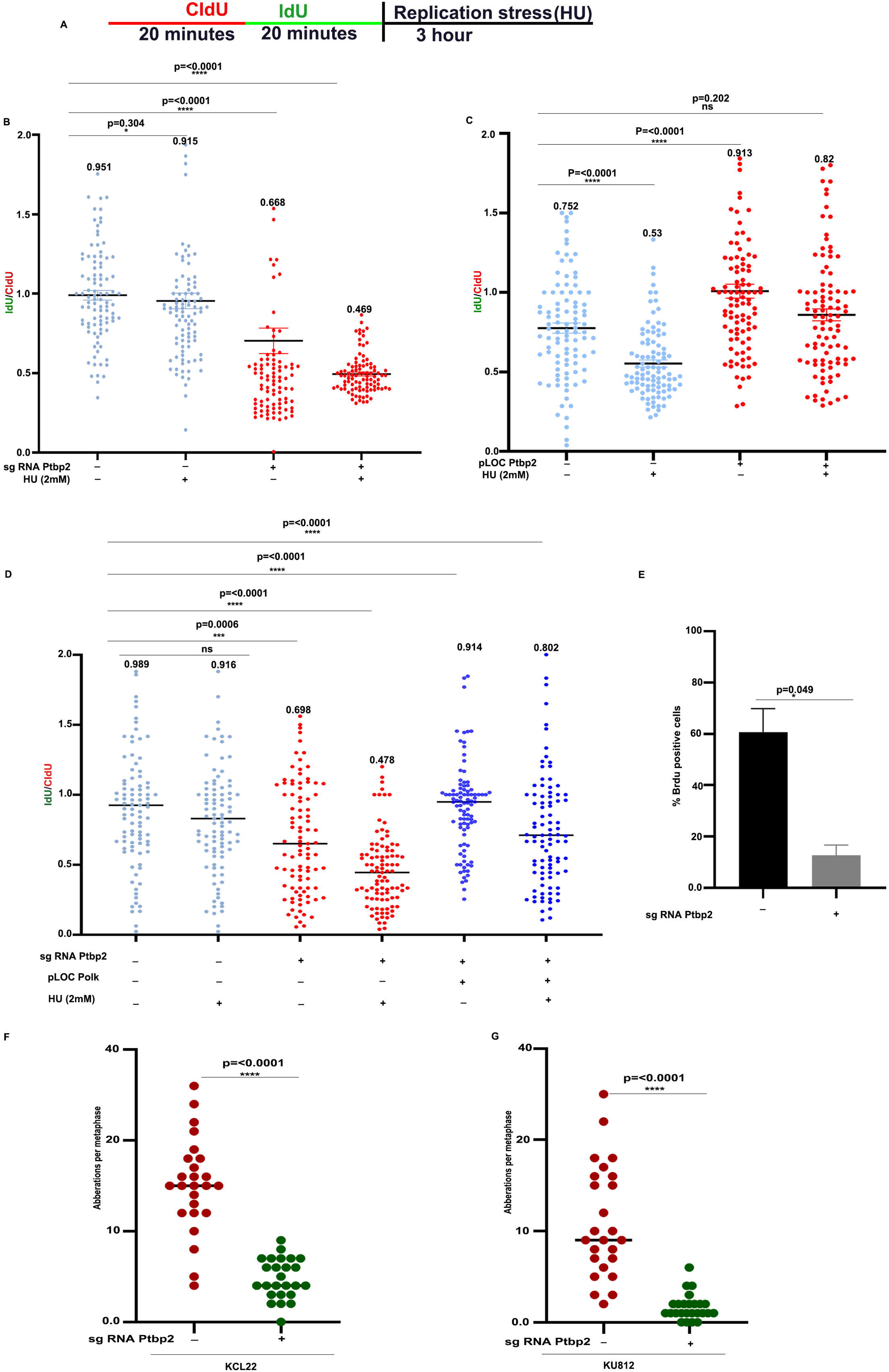
PTBP2 protects the stalled replication forks from degradation and promotes genomic instability. A) Schematic for labeling cells with CIdU and IdU. B) Ratios of IdU versus CIdU in the WT NTC, Ptbp2-KO-KCL22 cells treated with or without hydroxyurea. C) Ratio of IdU versus CIdU in the LAMA84 and LAMA84 Ptbp2 O/E cells with or without hydroxyurea treatment. (NS, not significant). One hundred replication forks were analyzed for each genotype. D) Ratio of IdU versus CIdU in the KCL22-NTC, Ptbp2-KO-KCL22, and Ptbp2-KO-Polκ-O/E-KCL22 cells treated with or without hydroxyurea. (NS, not significant). E) Quantification of the percentage of the Brdu-positive cells. (F and G) 25 metaphases were counted for each condition, and the number of SCEs was normalized to the number of chromosomes per metaphase, which is represented in the graph for KCL22 and KU812 cells, respectively.

Increased cellular proliferation and genomic instability are the hallmarks of cancer cells. PTBP2 has been previously shown to increase cellular proliferation (**Barik et al. 2024**). To determine the effect of PTBP2 in the S phase, asynchronous *Ptbp2* ablated cells and their non-targeted control were analyzed for bromodeoxyuridine (BrdU) incorporation. We found that the incorporation of the Brdu was higher in the non-targeted control cells than the *Ptbp2* ablated cells, thereby supporting that PTBP2 protects replication forks from degradation and promotes genomic instability (**Fig. 4E)**.

Chromosomal aberrations are the hallmark of tumorigenesis. In CML, it has been reported that the progression of CML CP to CML BC is associated with the increase in chromosomal aberration from 7% to 75% (**Asnafi et al., 2018**); thus, we assessed the role of PTBP2 in chromosomal aberrations via the sister chromatid exchange (SCE) in NTC and *Ptbp2* ablated KCL22 and KU812 cells. The SCE was significantly higher in the *Ptbp2* NTC cells than in the *Ptbp2* ablated KCL22 and KU812 cells (**Supplementary Fig. 3**). Aberrations per metaphase in *Ptbp2* ablated KCL22 and KU812 cells are shown in **Fig. 4F and 4G**, respectively.

### PTBP2 promotes tumor progression by increasing genomic instability

Recently, we reported that PTBP2 promotes tumor formation as mice injected subcutaneously with KCL22-NTC cells were associated with large tumors, and the *Ptbp2*-KO-KCL22 cells gave comparative smaller-sized tumors and were less aggressive, as shown by reduced Ki67 expression. In the tissue samples derived from the tumor of the control mice, cellular pleomorphism, a characteristic feature of malignancy, was more evident when compared to the tissue samples derived from the tumor of the mice with *Ptbp2*-KO-KCL22 cells (**Barik et al., 2024)**. PTBP2 and POLK were highly overexpressed in the KCL22-NTC tumors compared to the *Ptbp2*-KO-KCL22 tumors in both the mRNA and protein levels (**Fig. 5A and 5B**). The immunohistochemistry also confirmed the overexpression of *Pol*_κ_ in the KCL22-NTC tumors compared to the *Ptbp2*-KO-KCL22 tumors (**Fig. 5C**, **upper panel**). The expression of γH2AX was significantly higher in the *Ptbp2*-KO-KCL22 tumor than in the KCL22-NTC cells (**Fig. 5C, lower panel**), thus indicating the extent of the unrepaired DNA in the cells. Histopathological evaluation was conducted to evaluate the mitotic score and the number of typical and atypical mitotic figures in the tumor tissues derived from the KCL22 control and *Ptbp2*-KO-KCL22 group mice. In the tumor tissue samples derived from the control group mice, many mitotic figures indicating active karyokinesis or cell proliferation could be observed (**Fig. 5D, upper panel**). The number of mitotic figures per high-power field was around 10 times more than the tumors derived from the *Ptbp2* knockout group. This was obtained by counting the mitotic figures per 10 high power fields (HPF) and observing the morphologic features of mitotic figures. The number of mitotic figures counted in 10 high-power fields is shown in the bar diagram (**Fig. 5E**). Apart from the number of mitotic figures, the tumors derived from the control group demonstrated a more significant number of atypical mitotic figures. These were primarily tripolar or multipolar mitotic figures, with more than 2 chromosome clusters, appearing as 3 or more linear metaphase plates of chromosomes, respectively (**Fig. 5D, 2^nd^ panel**). Asymmetrical mitotic figures with unequal-sized metaphase plates were another atypical form observed. Atypical mitotic figures refer to a dysregulated and random assembly of nuclear materials within the dividing cells, which reflects genomic instability, telomere dysfunction, and aneuploidy. These mitotic figures, reported as morphologic signatures of underlying genomic instability, are accredited as a feature of malignancy and associated with poor prognosis in various cancers. Another essential feature in the control group was the presence of polyploid/ multinucleated giant cells (**Fig. 5D, 3^rd^ panel**). These cells with higher DNA content have been reported to have varied functions that favor tumor progression, resistance to therapy, and cancer relapse.

**Figure 5.**
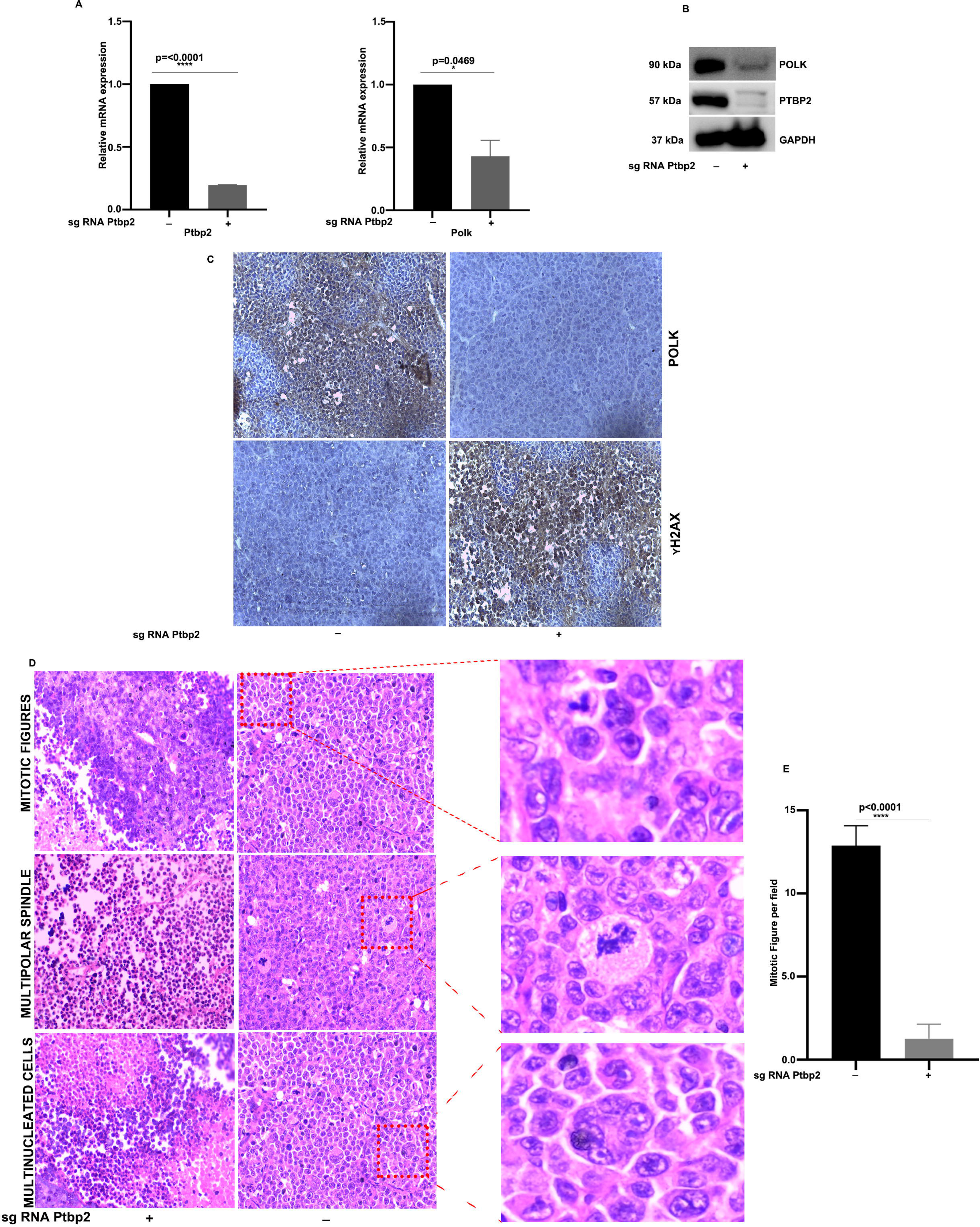
PTBP2 promotes tumor progression by increasing genomic instability. A) mRNA expression of Ptbp2 and Polκ in WT-NTC, Ptbp2-KO-KCL22 cell tumors B). Western blot analysis of PTBP2 and POLK from the protein isolated from KCL22-NTC, *Ptbp2*-KO-KCL22 cell tumors. GAPDH was used as a loading control. C). Representative images of IHC of tumor tissues stained with POLK and γH2AX antibody D). H&E staining of KCL22-NTC, *Ptbp2*-KO-KCL22 cell tumors. The 1st panel represents mitotic figures, the 2nd panel represents the multipolar spindle, and the 3rd panel represents the multinucleated cells. E). Quantification of the mitotic figures per field.

Again, we developed the BCR::ABL1 (CML) mice model by transducing the MSCV vector, BCR::ABL1, and BCR::ABL1+Ptbp2 in B6 CD 45.1 lineage-negative cells and transplanted the cells into the B6 CD 45.2 mice that were irradiated with 10Gy. After 30 days, the mice were sacrificed, and the spleen weight was checked and found to be higher for the BCR::ABL1 and BCR::ABL1+Ptbp2 cells (**Supplementary Fig. 4**). BCR::ABL1, *Ptbp2*, and *Pol*_κ_ mRNA expression were checked in the spleen. The expression of BCR::ABL1, *Ptbp2*, and *Pol*_κ_ was found to be increased, more so in the mice transplanted with BCR::ABL1+*Ptbp2* (**Supplementary Fig. 4**). Similarly, immunohistochemistry also suggested that the expression of POLK was higher in the BCR::ABL1+*Ptbp2* group for both the spleen and the liver than in control and BCR::ABL mice groups (**Fig. 6A and 6B).** Histopathological examination revealed standard histological architecture in the liver and spleen of the control group mice. The spleen of the BCR::ABL1 and the BCR::ABL1+*Ptbp2* group revealed effacement of the typical architecture with increased infiltration of neoplastic myeloid cells, leading to increased cellularity and a decrease in the proportion of the red pulp. The neoplastic cells were seen invading the splenic trabeculae and the capsule (**Fig. 6C, upper panel**). Incredibly abnormal mitotic figures and multinucleated cells were also observed. Such abnormal mitotic figures were significantly higher in the BCR-ABL+*Ptbp2* group, indicating increased genomic instability (**Fig. 6D**). The liver of these two groups showed infiltration of leukemic myeloid cells forming nodular collection in the sinusoids, especially in the portal area. The number and size of these nodules were comparatively more significant in the BCR-ABL1+*Ptbp2* group, indicating more aggressive behavior of these cells. The surrounding hepatocytes showed degenerative and necrotic changes, more pronounced in the BCR-ABL1+*Ptbp2* group (**Fig. 6C, lower panel).** Thus, it seems from both the mice models that the presence of PTBP2 and the expression of BCR::ABL1 facilitates the disease progression in CML cells by increasing genomic instability.

**Figure 6.**
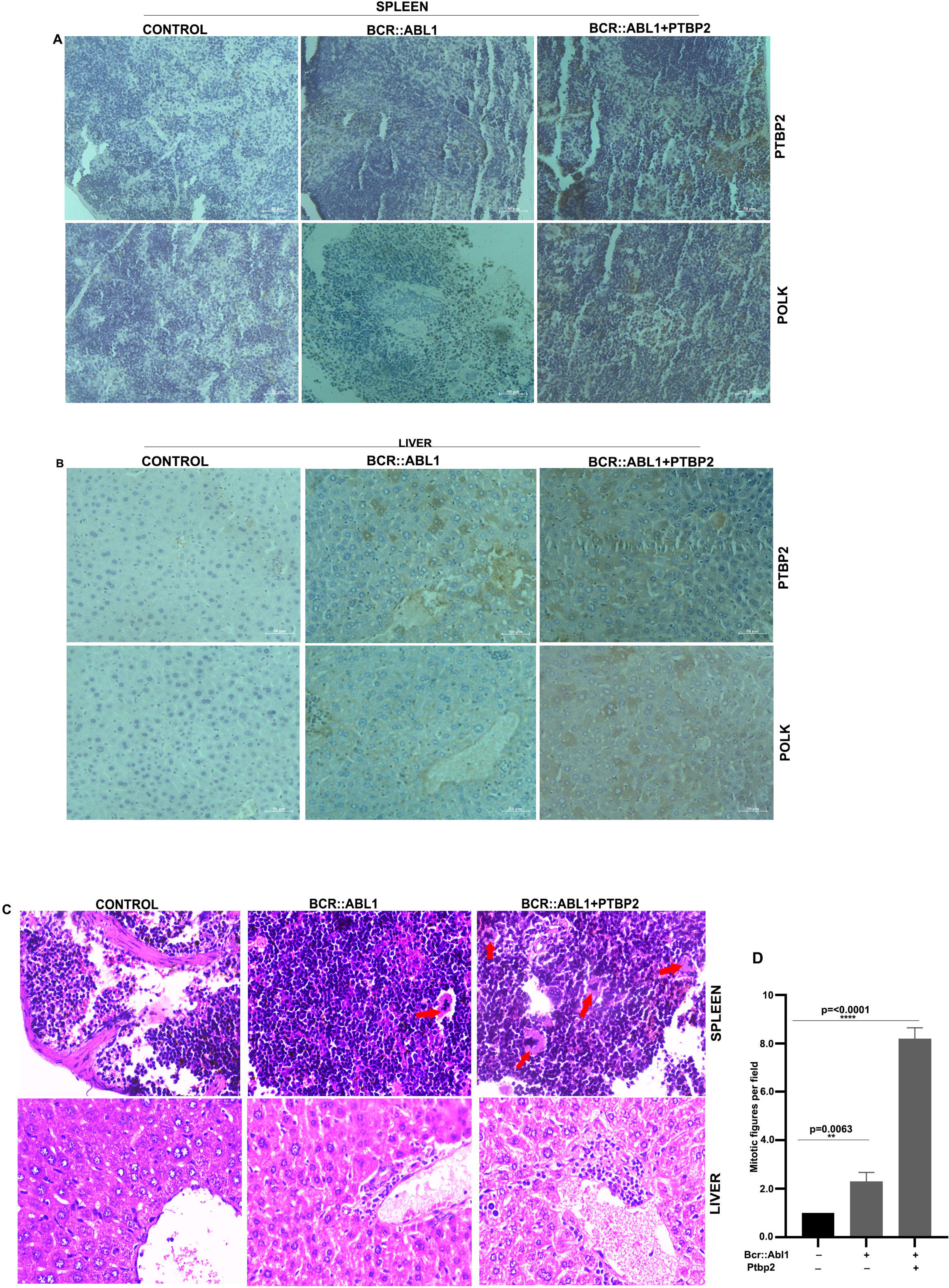
PTBP2 and BCR::ABL promote aggressive CML disease by increasing genomic instability. A and B) Representative images of IHC from the spleen and liver of the respective groups probed with POLK and PTBP2 antibodies. C) Liver-control mice-Normal histological arrangement-H&E X 400, Spleen-control mice-Normal histological arrangement-H&E X 400, BCR-ABL-Liver -neoplastic infiltration of myeloid cells-H&E X 400, BCR-ABL-Spleen - increased cellularity abnormal mitosis, - H&E X 400, BCR-ABL+PTBP2-Liver-neoplastic infiltration of myeloid cells-H&E X 400, BCR-ABL+PTBP2-Spleen-increased cellularity, abnormal mitosis, multinucleated cells -H&E X 400. D) Quantification of mitotic figures per field in BCR::ABL and BCR::ABL+PTBP2 transplanted mice.

## Discussion

Cancer development is driven by the accumulation of mutations, resulting in increased genomic instability and the ability of cancer cells to evade cell death. In CML, the BCR::ABL1 kinase has been found to trigger DNA damage by promoting the production of reactive oxygen species, thereby causing genomic instability (**Nowicki et al., 2004**). This instability is associated with resistance to imatinib treatment and the accumulation of chromosomal abnormalities (**Koptyra et al., 2006 and 2008**). Additionally, data suggests that the build-up of DNA damage, incorrect NHEJ repair processes, and changes in the DNA damage response play crucial roles in progressing to a more aggressive phase in CML patients (**Popp et al., 2020**). Moreover, the excessive production of white blood cells in CML can damage DNA as these cells undergo rapid division and proliferation, accumulating errors and mutations in their DNA. Similarly, specific translocations in acute myeloid leukemia, such as FLT3/ITD, and myelodysplastic syndrome, such as Ras mutation, have been shown to induce DNA damage and disrupt the regulation of critical repair proteins (**Sallmyra et al., 2008**). Eventually, these perturbations lead to the accumulation of enough mutations to promote the growth of neoplastic cells. Recently, we have shown the role of *Ptbp2* as an oncogene in CML (**Barik et al., 2024**). Presently, we show the importance of PTBP2 in cancer progression, as its expression promotes the upregulation of *Pol*_κ_. *Pol*_κ_, an error-prone DNA polymerase, is up-regulated in non-small cell lung carcinoma, melanoma, and glioblastoma, which induces DNA breaks, stimulates DNA exchanges, and promotes aneuploidy (**O-Wang et al., 2001; Temprine et al., 2020; Peng et al., 2016)**. Reduced fidelity of the replication machinery due to an overrepresentation of *Pol*_κ_ could accelerate tumorigenesis by giving a selective growth advantage during cancer cell evolution. Ablation of *Ptbp2* and, thereby, downregulation of *Pol*_κ_ lead to the enhancement of expression of H2AX phosphorylation and γ-H2AX foci formation in the cells, a hallmark signal for DNA damage. This was found to be further increased when the cells were treated with HU. Overexpression of *Pol*_κ_ in the *Ptbp2* KO cells blunted H2AX phosphorylation and γ-H2AX foci formation with HU treatment. Our data also provide evidence that the PTBP2-POLK axis favors cell proliferation, as KO of *Ptbp2* and subsequent treatment of cells with HU deregulated the replicative potential of the cells and promoted apoptosis. The MRE11/RAD50/NBS1 (MRN) complex is a crucial sensor that detects damaged DNA and recruits ATM to DNA foci for activation (**Lee and Paull, 2005; Paull and Lee, 2005**). The MRE11 nuclease plays critical functions at DSBs and at stalled, collapsed, or reversed replication forks (**Schlacher et al., 2011; Stracker and Petrini, 2011; Mijic et al., 2017**). As a complex with RAD50 and NBS1, it plays a significant role in detecting and endonucleolytic processing of DNA ends, activating the ATM kinase. Our data show upregulation of the MRN complex-related protein in the presence of PTBP2. Similarly, ATM-CHK2 was upregulated in the presence of *Ptbp2*, whereas the genetic ablation of *Ptbp2* resulted in the whole axis’s shutdown. It has been reported that activation of the ATM-CHK2 cascade is essential for cell survival as it triggers a series of downstream pathways critical for DNA repair. **Paull and Lee, 2005** have previously reported some exciting findings about the physical interaction of MRE11 that directly stimulates ATM kinase. Following this, we checked the regulation of the MRN complex via POLK, which depicted the regulation of MRE11 but not RAD50 and NBS1 by POLK. MRE11 interacted with POLK as the pulldown of the POLK coimmunoprecipitated MRE11, consistent with the reverse coimmunoprecipitation and the colocalization assay. The interaction between POLK and MRE11 appears to influence the expression of ATM-CHK2. Also, the overexpression of *Pol*_κ_ in *Ptbp2* KO cells restored the MRE11-ATM-CHK2 pathways. This confirms that PTBP2 regulates the pathway through *Pol*_κ_. Because of decreased accuracy in copying DNA, the error-prone DNA polymerases, which specialize in bypassing DNA damage, can lead to random mutations in regions of the genome that are not damaged. This fosters genomic instability, leading to the formation of aggressive tumors. As a result of these effects, an excess of POLK in the CML blast crisis, along with the BCR::ABL1, promotes more aggressive tumor formation in mice. Our animal studies demonstrate that excess PTBP2-POLK expression promotes several atypical mitotic figures and polyploid/multinucleated giant cells, leading to aggressive tumors. The data presented here suggests a possible mechanism wherein PTBP2 promotes genomic instability by regulating the POLK-MRE11-ATM-CHK2 axis, thereby opening a novel area for treatment. This study has clinical implications as recently, it was shown that POLK is crucial for tolerating cisplatin-induced DNA damage lesions. When *Pol*_κ_-deficient tumors are treated with cisplatin, the outgrowth of tumors is significantly delayed, and the overall survival of tumor-bearing mice is increased (**Spanjaard et al., 2022**). Again, *Pol*_κ_ plays a role in mediating homologous recombination repair and temozolomide resistance in glioblastoma through Rad17-dependent activation of ATR-CHK1 signaling and deregulation of *Pol*_κ_ sensitizes cells towards temozolomide (**Peng et al., 2016; Ribeiro et al., 2024**). Our previous findings have shown that suboptimal concentration of imatinib and knockout of *Ptbp2* effectively reduced the cell viability compared to imatinib alone (**Barik et al., 2024)**. Thus, efficient but unfaithful DNA repair and enhanced survival capacity of the cells in the CML blast crisis observed in this study may preclude the cells from the inhibitory effects of imatinib and may contribute to drug resistance. Our findings highlight the problem and provide potential therapeutic targets, wherein small molecule mediated depletion of PTBP2 might reduce the progression of the disease as the transformation to the blastic phase was associated with the overexpression of *Pol*_κ_ and *Ptbp2*, offering hope for developing effective CML blast crisis therapy strategies. Also, we speculate from our findings that this approach may work for all cancers where *Pol*_κ_ is upregulated.

## Acknowledgment

SL and BB are supported by a fellowship from the Council of Scientific and Industrial Research (CSIR), India. TS, SC, and MB are supported by a fellowship from the University Grants Commission (UGC), India. SL, BB, TS, SC, and MB are registered with the Regional Centre of Biotechnology, Faridabad, India. The pCMV PTBP2 plasmid was kindly provided by Dr. Miriam Llorian (University of Cambridge, Department of Biochemistry, Cambridge, UK). The authors would like to acknowledge the help of Dr. Payel Guha and S.N. Rajashree for some molecular and cell biology work. The authors would like to acknowledge the assistance of Mr. Bhabani Sahoo of the Confocal facility, Mr. Paritosh Nath of the Flow Cytometry facility, Dr. Sarita Jena, and Mr. Biswajit Patra of the Animal House facility.

## Author contribution

SL, BB, and SC* designed and performed the experiments. TS, SC¥, and MB performed RT-qPCR and cloning. SIS performed the H&E staining and interpretation. SM, SB, and GB provided clinical inputs. SC* arranged for funding, designed and supervised the experiments, interpreted the data, and wrote the manuscript. All authors reviewed and approved the final version of the manuscript. SC^¥^: Sayantan Chanda; SC*: Soumen Chakraborty.

## Data Access Statement

Research data supporting this publication are available on request.

## Ethical approval and consent to participate

All animal protocols were approved by the Institute of Life Sciences (ILS/IAEC-03-AH/05-14 and ILS/IAEC-159-AH/AUG-19) Animal Ethics Committee. The Institutional Human Ethical Committee approved the study (16/HEC/12), and peripheral blood samples were collected from patients after obtaining written informed consent. All procedures involving human participants were performed according to the ethical standards of the institutional research committee and with the 1964 Helsinki Declaration and its later amendments or comparable ethical standards.

