## Supplementary figures and images for "DNA polymerase kappa stabilized by PTBP2 interacts with MRE11 and promotes genomic instability in leukemia cells"

### Supplementary Fig. 1

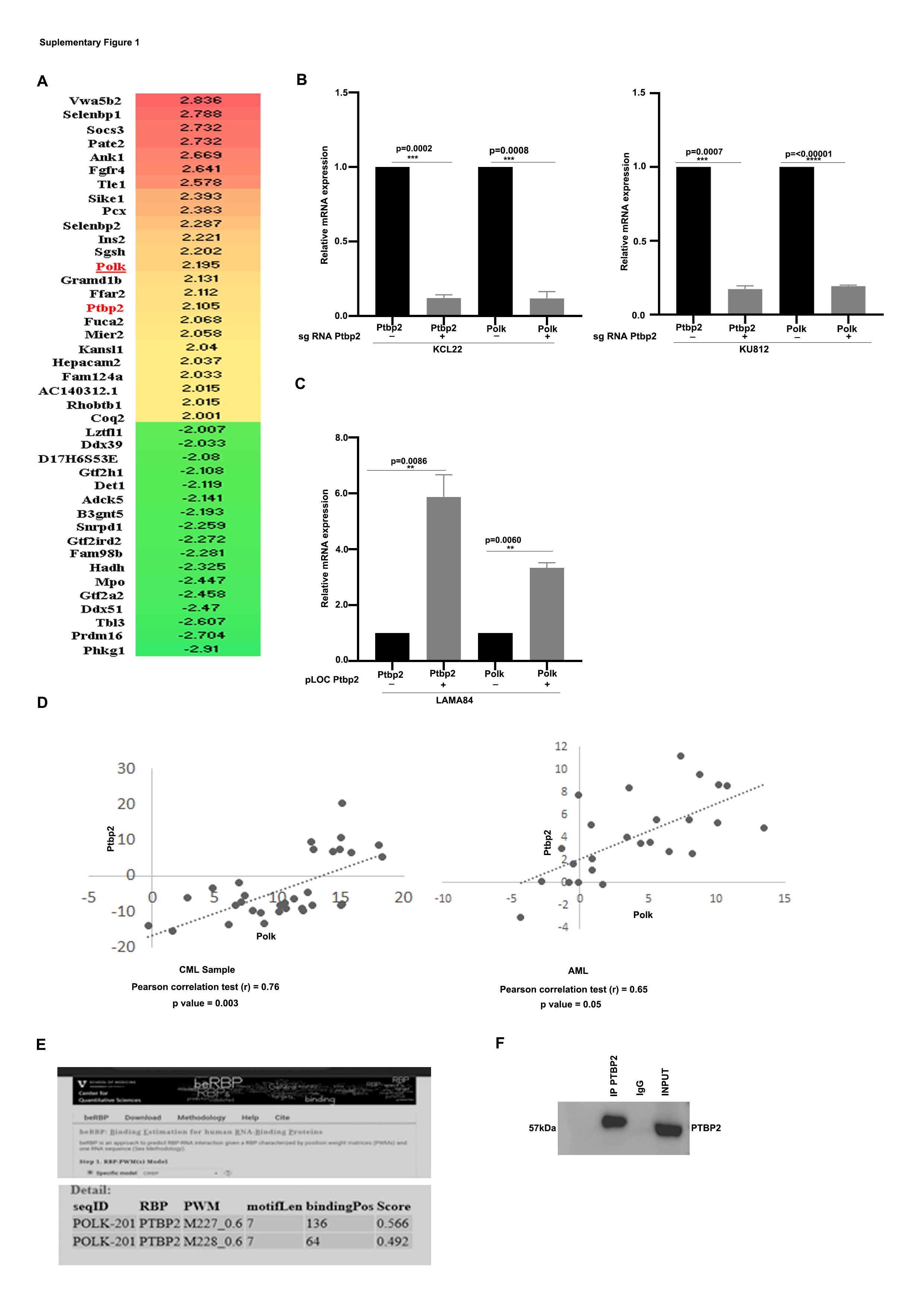

### Supplementary Fig. 2A

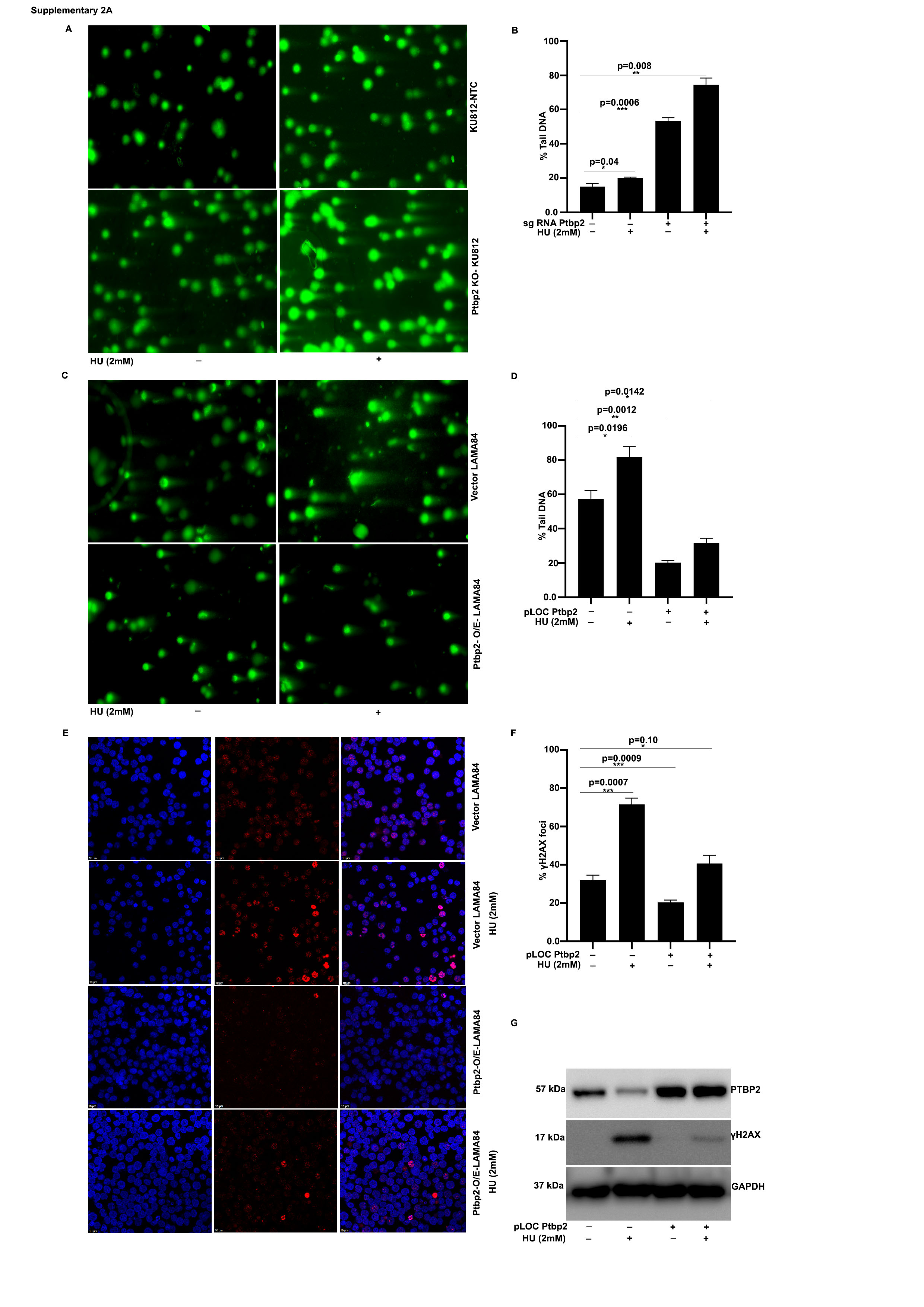

### Supplementary Fig. 2B

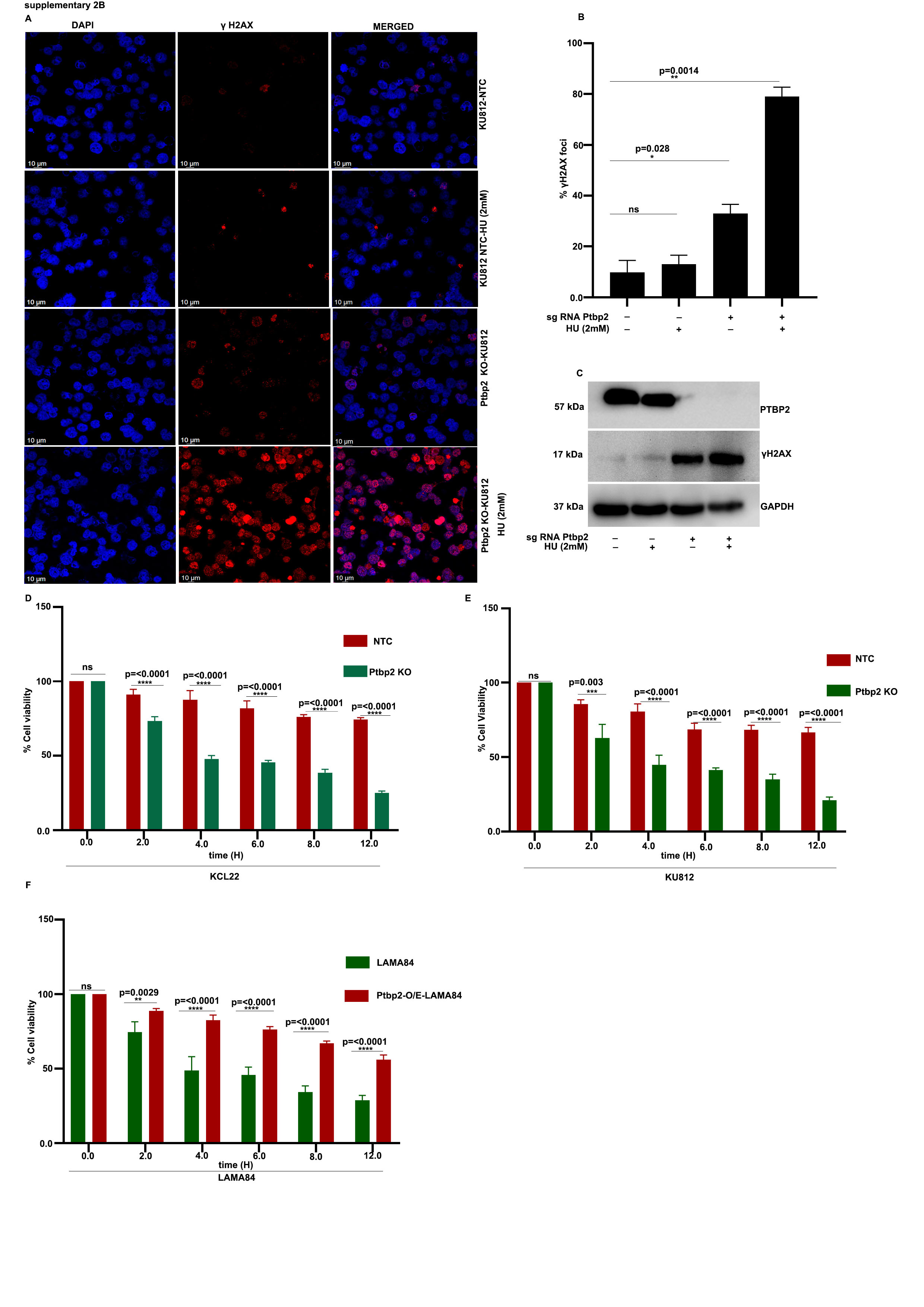

### Supplementary Fig. 2C

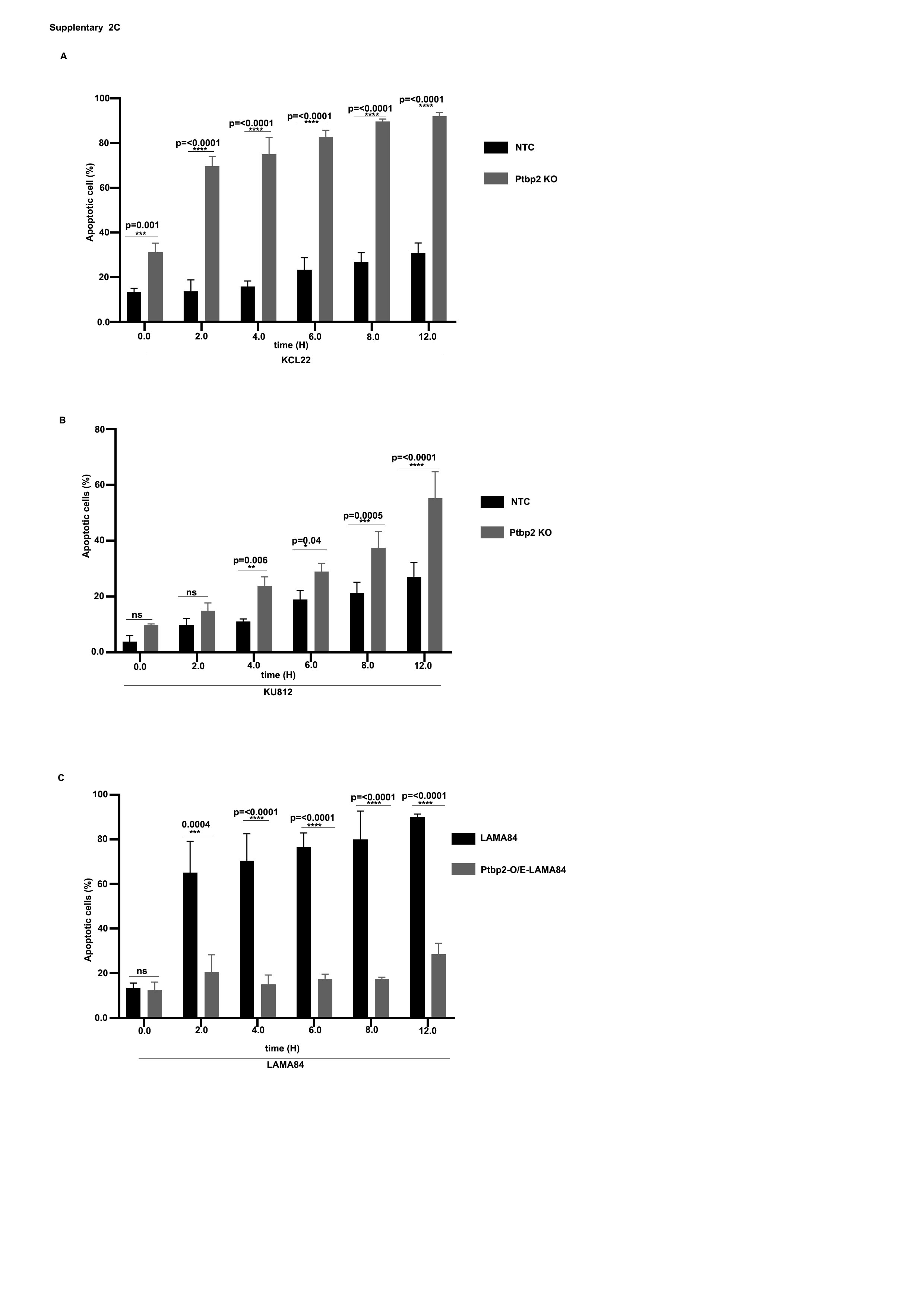

### Supplementary Fig. 3

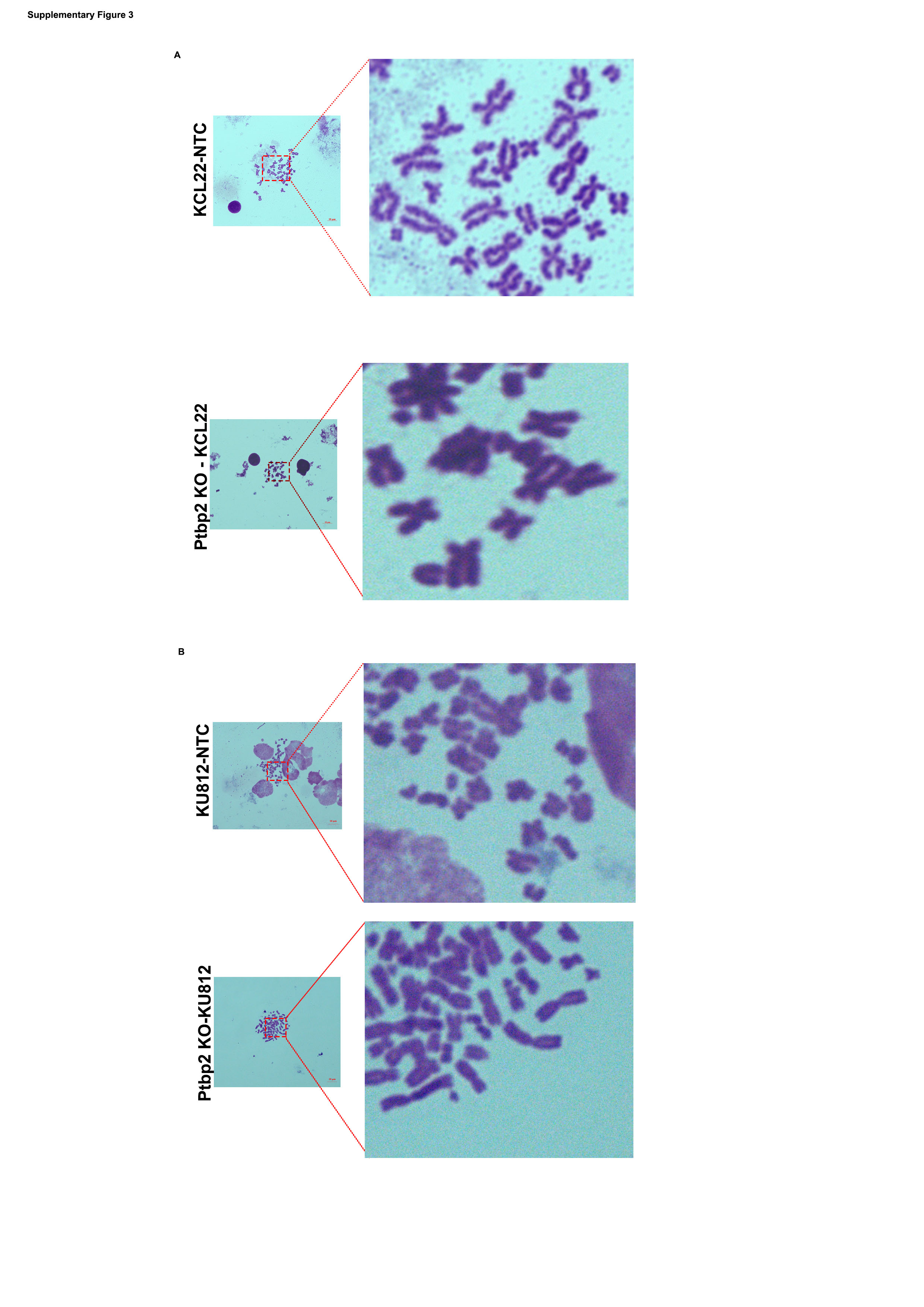

### Supplementary Fig. 4

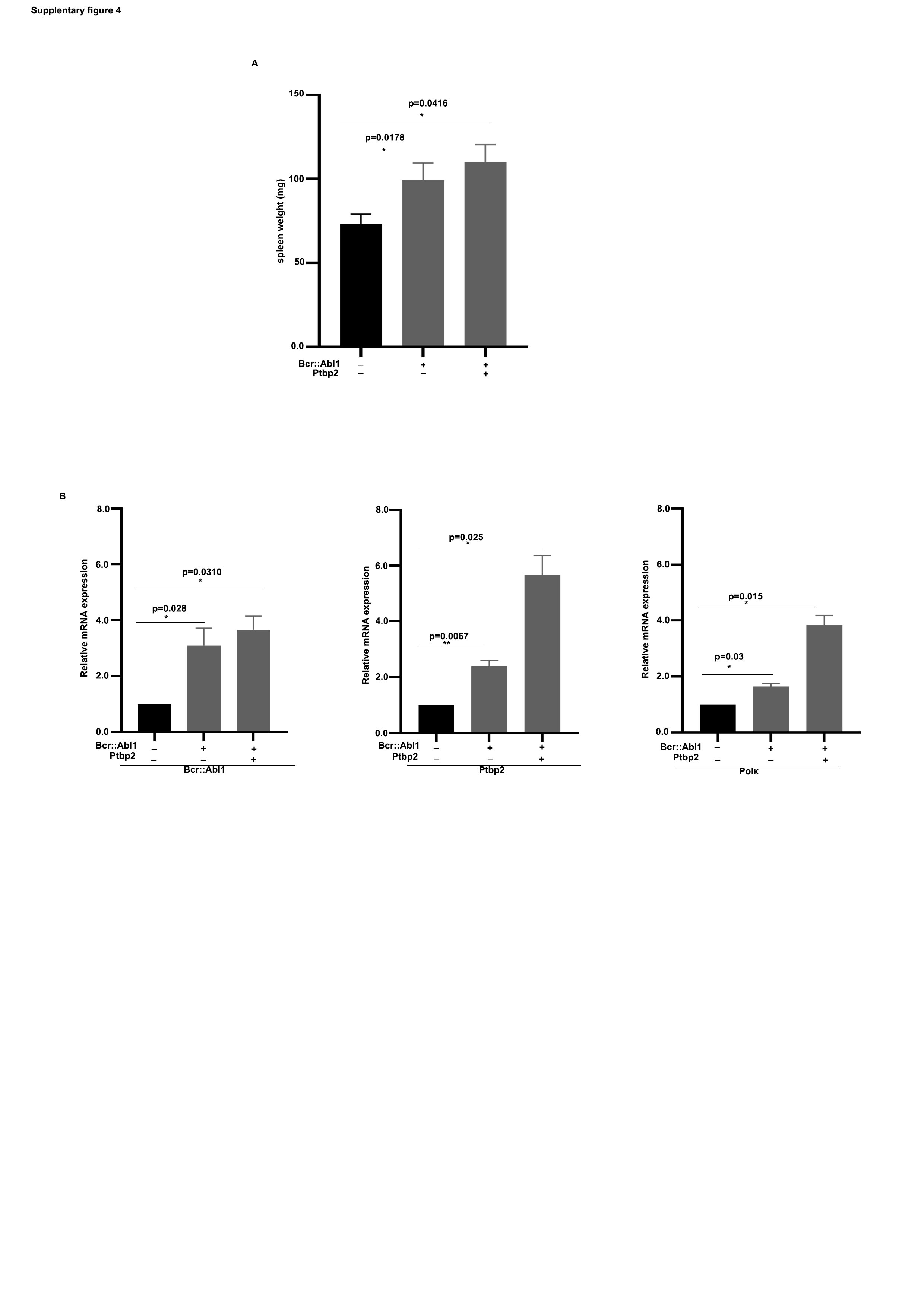
