## Supplementary Figure legends for "DNA polymerase kappa stabilized by PTBP2 interacts with MRE11 and promotes genomic instability in leukemia cells"

**Supplementary Figures**

Supplementary Figure 1 **PTBP2 targets DNA polymerase kappa and facilitates its upregulation.**

A) Heatmap of the differentially expressed genes in vector-32Dcl3 compared to 32Dcl3-Ptbp2 cells. The average log_2_ fold from two 32Dcl3-Ptbp2 cells is shown. B) mRNA expression of Ptbp2 and Polκ in Ptbp2 NTC and Ptbp2 ^-/-^ cells by RT-qPCR in KCL22 and KU812 cells. C) mRNA expression of Ptbp2 and Polκ in Ptbp2 overexpressed LAMA84 cells. D) Correlation of Ptbp2 and Polκ is achieved in different CML (n=34) and AML samples (n=34). E) Predicting the binding site of Ptbp2 in the 3’UTR of Polk using the publicly available database beRBP. F) The western blot checked the Immunoprecipitation of Ptbp2 and the input control.

Supplementary Figure 2	**PTBP2-POLK axis acts as a regulator of DNA repair in hematological malignancies.**

A) KU812-NTC and Ptbp2-KO-KU812 cells treated with or without hydroxyurea and subjected to alkaline comet assay were performed. B) Quantification of the percentage of DNA tail. The data are presented as the mean ± SEM. C) LAMA84 and LAMA84 O/E Ptbp2 cells treated with or without hydroxyurea and subjected to alkaline comet assay were performed. D) Quantification of the percentage of DNA tail. The data are presented as the mean ± SEM. E) LAMA84-vector and Ptbp2-O/E-LAMA84 cells treated with or without hydroxyurea and probed with γH2AX antibody. F) Quantitative analysis of the percentage of γH2AX foci. A cell containing at least 10 foci was considered a foci-positive cell. G) Protein expression of γH2AX and PTBP2 in LAMA84 and LAMA84 O/E Ptbp2 cells treated with or without hydroxyurea.

Supplementary 2B

A) KU812 NTC, Ptbp2-KO-KU812 cells treated with or without hydroxyurea and stained with γH2AX antibody. B) Quantitative analysis of the percentage of γH2AX foci. A cell containing at least 10 foci was considered a foci-positive cell. C) Protein expression of γH2AX and PTBP2 in KU812 NTC, Ptbp2-KO-KU812 treated with or without hydroxyurea.

D) Cell viability of KCL22- NTC, Ptbp2-KO cells, and vector-LAMA84 & Ptbp2-O/E-LAMA84 cells treated with or without hydroxyurea for different time points (2,4,6,8 and 12 h respectively). The graph represents the percentage of cell viability. The results were expressed as mean ± SEM of triplicate experimental analysis.

Supplementary 2C

**A)** KCL22- NTC, Ptbp2-KO cells, KU812-NTC, Ptbp2-Ko cells, and vector-LAMA84 & Ptbp2-O/E-LAMA84 cells treated with or without hydroxyurea for different time points (2,4,6,8 and 12 h respectively). The graph represents the percentage of Apoptotic cells. The results were expressed as mean ± SEM of triplicate experimental analysis.

Supplementary Figure 3 **PTBP2 protects the stalled replication forks from degradation and promotes genomic instability.**

A) Representative images of SCEs Giemsa-stained chromosome spreads from WT-NTC, Ptbp2-KO-KCL22 cells, and Ptbp2-KO-KU812 cells.

Supplementary Figure 4 **PTBP2 and BCR::ABL promote aggressive CML disease by increasing genomic instability.**

A) Spleen weight of mice transplanted with MSCV vector BCR::ABL and the BCR::ABL+Ptbp2 transduced cells. B) Relative mRNA expression of Bcr::Abl, Ptbp2, and Polκ in mice transplanted with MSCV vector, BCR::ABL, and the BCR::ABL+Ptbp2 transduced cells.
