## Supplementary Table for "DNA polymerase kappa stabilized by PTBP2 interacts with MRE11 and promotes genomic instability in leukemia cells"

| Sl. No | Primer | Sequence |
| --- | --- | --- |
| 1 | PTBP2-F | TCTGAGTCCTTTGGCCATTC |
| 2 | PTBP2-R | GGTAAACAGACTTTGGGGCGTA |
| 3 | H GAPDH-F | ACCCACTCCTCCACCTTTG |
| 4 | H GAPDH-R | CTCTTGTGCTCTTGCTGGG |
| 5 | M GAPDH-F | AATCCCATCACCATCTTCCA |
| 6 | M GAPDH-R | CCAGGGTCTTACTCCTTG |
| 7 | H POLK-F | GCACAAAGGAGAAGTGTGACAG |
| 8 | H POLK-R | CCCCTTCGTGGCTTCCATTA |
| 9 | M POLK-F | TCACCTGCTGCCATAGCTAATCT |
| 10 | M POLK-R | GGCGATGCCTTAAAAAGAAGAG |
| 11 | M VWA5b2-F | CCCTGGAGTTTATGAGGTGGCA |
| 12 | M VWA5b2-R | GGAAGTAAGCCTGTCCTCTGCT |
| 13 | M Selenbp1-F | CTGATACTGCCTGGTCTCA |
| 14 | M Selenbp1-R | AGTGGCTGGTGTGCAAAC |
| 11 | M Pate2-F | CATGACATCCTGCGTACCAAATC |
| 12 | M Pate2-R | GATGTCCTCACAGTTGGTCATAC |
| 13 | M Tle1-F | GAACGTGCCAAACAGGTGAC |
| 14 | M Tle1-R | TGTTGATCTGCCGAGCATGT |
| 15 | M Sike1-F | AGCAGTGATGCGGAGAGCTGTT |
| 16 | M Sike1-R | GCAGAGATTCACTGCTGATGGAC |
| 17 | M Pcx-F | GGATGACCTCACAGCCAAGCAT |
| 18 | M Pcx-R | GCAATCGAAGGCTGCGTACAGT |
| 19 | M Selenbp2-F | CTGATACTGCCTGGTCTCA |
| 20 | M Selenbp2-R | AGTGGCTGGTGTGCGTAT |
| 21 | M Ins2-F | CGTGGCTTCTTCTACACACCCA |
| 22 | M Ins2-R | TCCAGTGCCAAGGTCTGAAGGT |
| 23 | M Sgsh-F | TACGTGGCTTTCCATGACCC |
| 24 | M Sgsh-R | CTAGAAGGCTCACGTAGGCG |
| 25 | M Gramd1b-F | TACGTGGCTTTCCATGACCC |
| 26 | M Gramd1b-R | CTAGAAGGCTCACGTAGGCG |
| 27 | M Ffar2-F | CCACTGTATGGAGTGATCGCTG |
| 28 | M Ffar2-R | GGGTGAAGTTCTCGTAGCAGGT |
| 29 | M Fuca2-F | GGAACAGCACTGGCTTCTTAGC |
| 30 | M Fuca2-R | GCCACCATGTTTGCAGATAGACC |
| 31 | M Rhobt1-F | CTGTCAGTGGTGTGAGCATCGA |
| 32 | M Rhobt1-R | CAGACGCTGTTGTAGTTGGTGC |
| 33 | M Coq2-F | GCGTCCTTACTTCTTGTCGTCAC |
| 34 | M Coq2-R | CTTGACAGCAGACCATCCGAGT |
| 35 | M Lztfl1-F | TGTCCCAAATACGCCTCTGT |
| 36 | M Lztfl1-R | GGATGGCTCTTCTTCCCACT |
| 37 | M DDX51-F | CCCTGGTTACGGGACAGAAG |
| 38 | M DDX51-R | GTCAATCATCCGGTCAGCCT |
| 39 | M Kansl1-F | TGTTGTCCCGCTCAAGAGTC |
| 40 | M Kansl1-R | TAGAGGGAAGGGAAGAGCCC |
| 41 | POLK 3’UTR F1 site2 | GTTGGCATTGATCCTAGTG |
| 42 | POLK3’UTR Reverse | ACTGATTCAAGTGCTTATTC |
| 43 | POLK 3’UTR M-F | TTTTAATTTGGCAAAACATATTTAAAATTC |
